# GOALS: Gene Ontology Analysis with Layered Shells for Enhanced Functional Insight and Visualization

**DOI:** 10.1101/2025.04.22.650095

**Authors:** Zongliang Yue, Robert S. Welner, Christopher D. Willey, Rajesh Amin, Jake Y. Chen

## Abstract

Gene Ontologies (GOs) are standardized descriptions of gene functions in terms of biological processes, molecular functions, and cellular components, capturing their Parent-Child relationships in a structured framework and advancing cancer biological modeling to provide consistent and meaningful insights into functional genomics analysis. The conventional GO hierarchical structure is defined by human curation experts, with levels determined by the shortest path to the root term. However, grouping GOs poses challenges due to the uneven distribution of gene members within GO terms and inconsistencies in the level of detail across terms at the same GO level.

In this work, we introduce Gene Ontology Analysis using Layered Shells (GOALS), a novel tool that discretizes GOAs into optimal layers. GOALS creates scalable GO layers while maintaining a balanced number of genes across GOs in each layer. Unlike existing tools, the GOALS framework organizes GO terms using a bottom-up approach based on their co-membership network, discretizing GOs to achieve an exponential fit with GO’s gene member size. Meanwhile, GOALS reveals clusters or supersets reflecting biological relevance by unsupervised clustering of GO’s latent projections.

In a case study on mouse natural killer (NK) cell development, GOALS identified distinct GO functional clusters with multi-GO layers to reveal multiple levels of detail from specific to abstract contexts to maximize signal discovery and uncover those signals’ associations with trajectory divergence. More importantly, GOALS enhances enrichment analysis by introducing additional GO stratification and latent GO map that enables more accurate classification of functional differences.

GOALS offers a robust and innovative framework for exploring disordered GO clusters, mining GO activities, and analyzing potential GO-GO interplays. By addressing critical challenges in functional genomics, GOALS provides a powerful tool for advancing our understanding of cell heterogeneity and potentially uncovering actionable insights for therapeutic development.

## Introduction

Gene Ontology Annotation (GOA) plays a vital role in functional genomics by providing standardized and comprehensive annotations using Gene Ontology (GO) terms(1-3). To enhance interpretability, GOA incorporates hierarchical “is-a” relationships, enabling the capture of different levels of specificity. In this structure, child terms denote more specific biological functions than their parent terms. Unlike a strict tree hierarchy, a single term has multiple parent terms, reflecting the complex and interconnected nature of biological processes. This flexible architecture allows users to systematically reason through biological categories and their subtypes, offering a flexible “zoom in” and “zoom out” perspective, from high-level concepts to fine-grained details that are often overlooked by flat visualizations(4-6). Such hierarchical reasoning is especially essential in comparative genomics, where understanding functional differences at various levels of abstraction goes beyond pathway analyses(7-9). In functional enrichment analysis, this structure allows researchers to either roll up to broader GO categories or drill down to more specific terms, facilitating the identification of genes involved in general or specific functions. Moreover, the scalable design of GOA supports the integration of new GO terms without disrupting existing definitions, ensuring adaptability as biological knowledge expands. Currently, GOA encompasses three major ontological domains, including Biological Processes (GO-BP), Molecular Functions (GO-MF), and Cellular Components (GO-CC).

Scaling GO terms can be challenging, especially when multiple trends of GO terms or functional groups have been involved(10). Since its inception in 1998, Gene Ontology Annotation (GOA) has grown nearly 9-fold from approximately 4,500 terms to 40,214 by March 2025, reflecting the continuous expansion of biological knowledge and the growing need to define functional specificity within its hierarchical “is-a” structure. The relationship in this structure is a directed acyclic graph (DAG), where GO terms have multiple parent terms that may reside at different levels of specificity based on the definitions and the number of associated gene members(11). This polyhierarchical structure adds complexity, making it challenging to isolate specific trends or understand how multiple trends relate without a comprehensive view of their relationships and advanced methods for reasoning through their distinctions. Meanwhile, the varying levels of granularity from broad categories to extremely specific subfunctions make it difficult to stratify and compare GO terms, even when they exist within the same network(12). Conceptual distinctions among GO terms can also lead to vague or ambiguous interpretations, particularly when their context-dependent relationships are not carefully considered(13).

The current GOA stratification approaches are insufficient for accurately capturing GO term specificity. Two major bottlenecks hindering effective stratification are inconsistent resolution and an excessive number of strata. Traditional methods, such as using GO level (based on the shortest path to the root)(14) or GO depth (based on the longest path to the root)(15), often result in high variability in gene set sizes within each stratum. This is primarily due to the design of the hierarchical structure, which reflects conceptual relationships between terms but does not account for the actual number of gene members annotated to each term. Alternatively, k-shell decomposition(16), though not inherently suited for hierarchical data, has been applied using a GO-to-GO co-membership network (m-type network), where GO terms are stratified into shells based on their topological location in the network. While this method reflects certain degrees of functional specificity, it produces more than 100 shells, complicating both interpretation and visualization. Therefore, a more consistent and interpretable stratification strategy is essential to enhance downstream analysis and biological insights. This includes the development of standardized metrics within a scalable multi-layer structure, designed to simultaneously address quality control, minimize selection bias, and prevent the loss of biologically relevant GO term hits(14).

In this paper, we introduce a novel framework called Gene Ontology Analysis using Layered Shell (GOALS), layer with significant improvements in GO term stratification, providing enhanced clarity, better biological interpretation, and improved visualization by mining their similarity in latent space. Specifically, GOALS introduces a scalable algorithm that discretizes GO terms into a manageable number of strata named GO layer, maintaining a balanced GO term gene size in each layer. It also improves consistency and visualization by resolving inconsistencies in GO gene size and reducing excessive shells compared to traditional k-shell decomposition-based methods. Furthermore, GOALS provides the GO map, a gene co-membership GO-to-GO network embedded in a latent space, featuring systematic GO term ranking through newly designed GO-level signal scores within a hierarchical structure. The GO map enhances functional clustering and interpretation through latent embeddings. The GO layer can be easily integrated into existing platforms or analysis tools, such as PAGER(17-19), AmiGO(20), GOrilla(21), g:profiler(22), and REVIGO(23). We demonstrate the effectiveness of our approach in a natural killer (NK) cell functional genomic study, successfully identifying distinct GO functional clusters within the multi-layer GO framework. This allows users to visualize biological signals from specific to abstract contexts.

## Results

### GO layer stratification via IAER using a scalable approach from specific to abstract contexts

The GO layer stratification formed by the IAER (Initiation, Assignment, Expansion, and Repetition) procedure introduces a scalable framework that organizes GO terms from specific to more abstract biological contexts. This four-step approach facilitates the construction of biologically coherent GO layers for downstream analysis (**Fig. 1**). In this framework, each node represents a GO term, and edges correspond to m-type GO-to-GO evaluated by the statistical significance test in PAGER(17). The initial shell numbers of a GO term are denoted by the first letter of its name (e.g., A, B, C) using k-shell decomposition(16), while node colors indicate their final layer assignments as determined by the GOALS framework.

**Figure 1.**
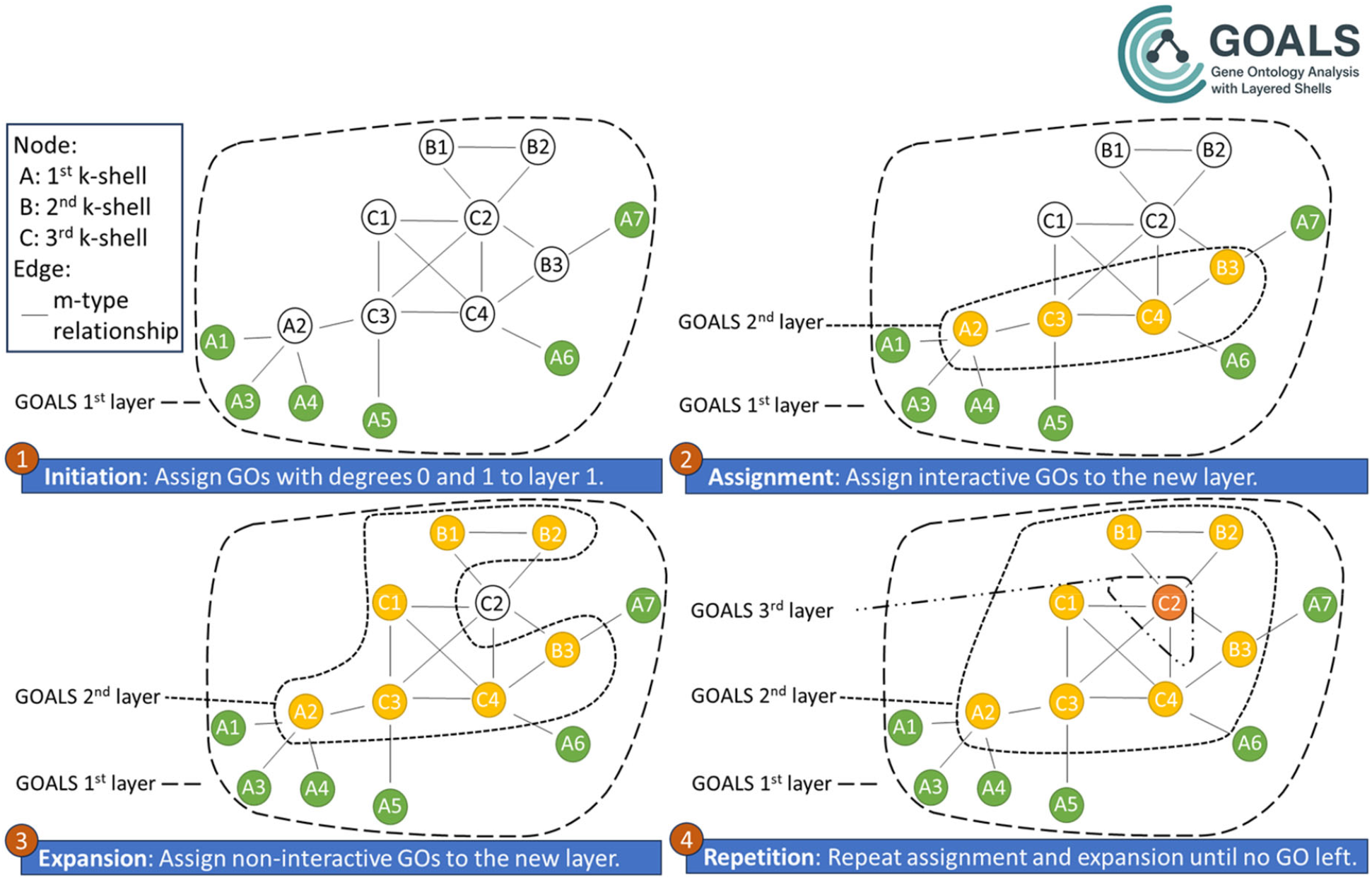
An example showcasing the four-step IAER procedure for generating stratified GO layers: Initiation, Assignment, Expansion, and Repetition. Nodes represent GO terms, while edges denote m-type GO-GO relationships. The initial shell number assigned to each GO is indicated by the first letter of its name. Node colors correspond to their GOALS layer assignment.

In the initiation step, GO terms with degrees 0 or 1 are identified and assigned to the first layer, as they exhibited minimal connectivity.

In the assignment step, GO terms that are connected via the m-type relationship to the current layer are identified and assigned to the next layer.

In the expansion step, non-interactive GO terms, which are not directly connected to previously layered nodes, are subsequently added to the new layer to maintain continuity and inclusion. A scalable threshold (*α*) is designed to filter the GO terms using the degrees of connections in the m-type network (see Methods).

In the repetition step, steps 2 and 3 are repeated iteratively until all GO terms have been assigned to a layer, ensuring a complete and non-redundant stratification across the entire GO network. This stratification approach enhances interpretability by structuring GO terms into biologically coherent layers, balancing GO size distribution while preserving functional connectivity.

The overall GOALS framework integrates four types of data: (1) the most up-to-date GO annotations and gene memberships, (2) GO depth and level information derived from the “is-a” hierarchical relationships, (3) differentially expressed genes (DEGs) identified through statistical analysis, and (4) phenotypic data from genomic studies (**Fig. S1**).

### GO m-type network optimization maximizing the power law distribution and GO term coverage in network

To optimize the m-type network, we perform a screening to find the best threshold for truncating the low confidence m-type relationship using the -log(PMF) value in maximizing the AI-adjusted R^2^ Index (ARI) in **Table 1** and **Table S1**. The coefficient of determination is derived from a linear regression between the logarithm transformed GO term degrees in the network and their log-transformed frequencies. Notably, as AI decreases, 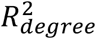 generally increases, indicating an inverse relationship between 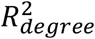 and AI (**Fig. S2** and **Fig. S3**). Therefore, by multiplying 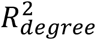 with the AI, we can identify the optimal - log(PMF) threshold that balances GO term coverage with a strong fit to the power-law distribution based on GO term degrees, ensuring high-quality network construction. In the GO-BP categories, when the - log(PMF) is set to be 7, GO-BP achieves nearly 100% term coverage in the m-type network with 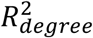 equals to 0.89, resulting in the highest IRI value of 0.891 (**Table 1**). In the GO-MF category, when the - log(PMF) is set to be 4, GO-MF has almost 100% term coverage in the m-type network with 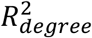 equals to 0.86, resulting in the highest IRI value of 0.857. In the GO-CC category, when the -log(PMF) is set to be 6, GO-CC has almost 99% term coverage in the m-type network with 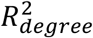 equals to 0.89, resulting in the highest IRI value of 0.878.

**Table 1.** The optimized -log(PMF) cutoff maximizing the AI-adjusted *R*^2^ index (ARI) of the *R*^2^ value of power-law distribution fit and the Aggregation Index (AI) of the m-type network in different GO categories.

| Category | -log(PMF) | ARI | $R^2_{degree}$ | AI |
| --- | --- | --- | --- | --- |
| BP | 7 | 0.891 | 0.89 | 1.00 |
| MF | 4 | 0.857 | 0.86 | 1.00 |
| CC | 6 | 0.878 | 0.89 | 0.99 |

### GO layer presents a novel GO term stratification with enhanced GO terms’ gene size consistency and significant reduction of variance using the mean of sample variance (MSV)

To optimize the GO layer stratification, we screen the optimal α-th percentile to find the best threshold for generating the scalable numbers of layers in maximizing the baseline-match-adjusted *R*^2^ index (BRI) in **Table 2** and **Table S2**. The coefficient of determination is derived from a linear regression between the logarithm-transformed GO frequency in the network and the total number of distinct GO layers. The baseline match score exhibits an elbow in the curve, which helps identify the optimal α threshold based on the baseline number of GO strata derived from GO depth and GO level (**Fig. S4**). In the GO-BP category, when the α is set to be 0.7, GO-BP achieves a baseline match score of 0.93 with 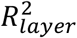 equals to 0.98, resulting in the highest BRI value of 0.912 (**Table 2**). In the GO-MF category, when the α is set to be 0.7, GO-MF achieves a baseline match score of 1 with 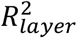 equals to 0.88, resulting in the highest BRI value of 0.88. In the GO-CC category, when the α is set to be 0.7, GO-CC achieves a baseline match score of 1 with 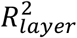 equals to 0.97, resulting in the highest BRI value of 0.97.

**Table 2.** The optimized α cutoff maximizing the product of the 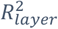 of power law distribution fit and the match score of stratum numbers in GO categories.

| Category | $\alpha$ | BRI | $R_{layer}^2$ | Baseline-match |
| --- | --- | --- | --- | --- |
| BP | 0.7 | 0.912 | 0.98 | 0.93 |
| MF | 0.7 | 0.880 | 0.88 | 1.00 |
| CC | 0.7 | 0.970 | 0.97 | 1.00 |

The GO layer outperforms traditional GO depth and GO level stratifications in capturing gene size consistency within each stratum and significantly enhances the fit to a power-law distribution, thereby improving biological interpretability (**Fig. 2**). Across all three GO categories, the GO layer shows substantial improvements in both the power-law fit 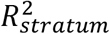 (**Fig. 2a**) and the reduction of the Mean of Sample Variance (MSV) (**Fig. 2b**). In the GO-BP category, the GO layer achieves a 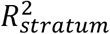 of 0.98 and a MSV of 1×10^4^, markedly better than the GO depth (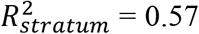 and MSV = 3×10^5^) and GO level (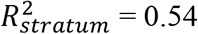 and MSV = 3×10^5^). In the GO-MF category, the GO layer yields a 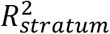 of 0.88 and MSV of 3×10^4^, compared with the GO depth (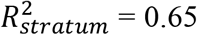 and MSV = 9×10^4^) and GO level of 0.62 and MSV of 9×10^4^. In the GO-CC category, the GO layer produces 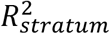 of 0.97 and MSV of 3×10^5^ representing strong improvement over the GO depth (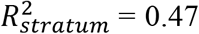 and MSV = 1×10^7^) and GO level (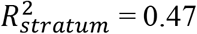 and MSV = 1×10^7^). The relationship between GO strata and the median gene size of GO terms within each stratum is well captured by a linear regression model, demonstrating a near-perfect fit to a power-law distribution, as illustrated in the boxplot (**Fig. 2c**). Additionally, the distribution of GO terms across strata shows that both GO depth and GO level tend to concentrate GO frequency in the middle stratum (neither specific nor abstract) across all three categories. In contrast, the GO layer redistributes GO term frequencies toward more specific stratum, resulting in significantly skewed distributions with skewness values of 1.43 in the GO-BP category, 1.74 in the GO-MF category, and 2.94 in the GO-CC category (**Fig. 2b**), indicating a more biologically meaningful stratification.

**Figure 2.**
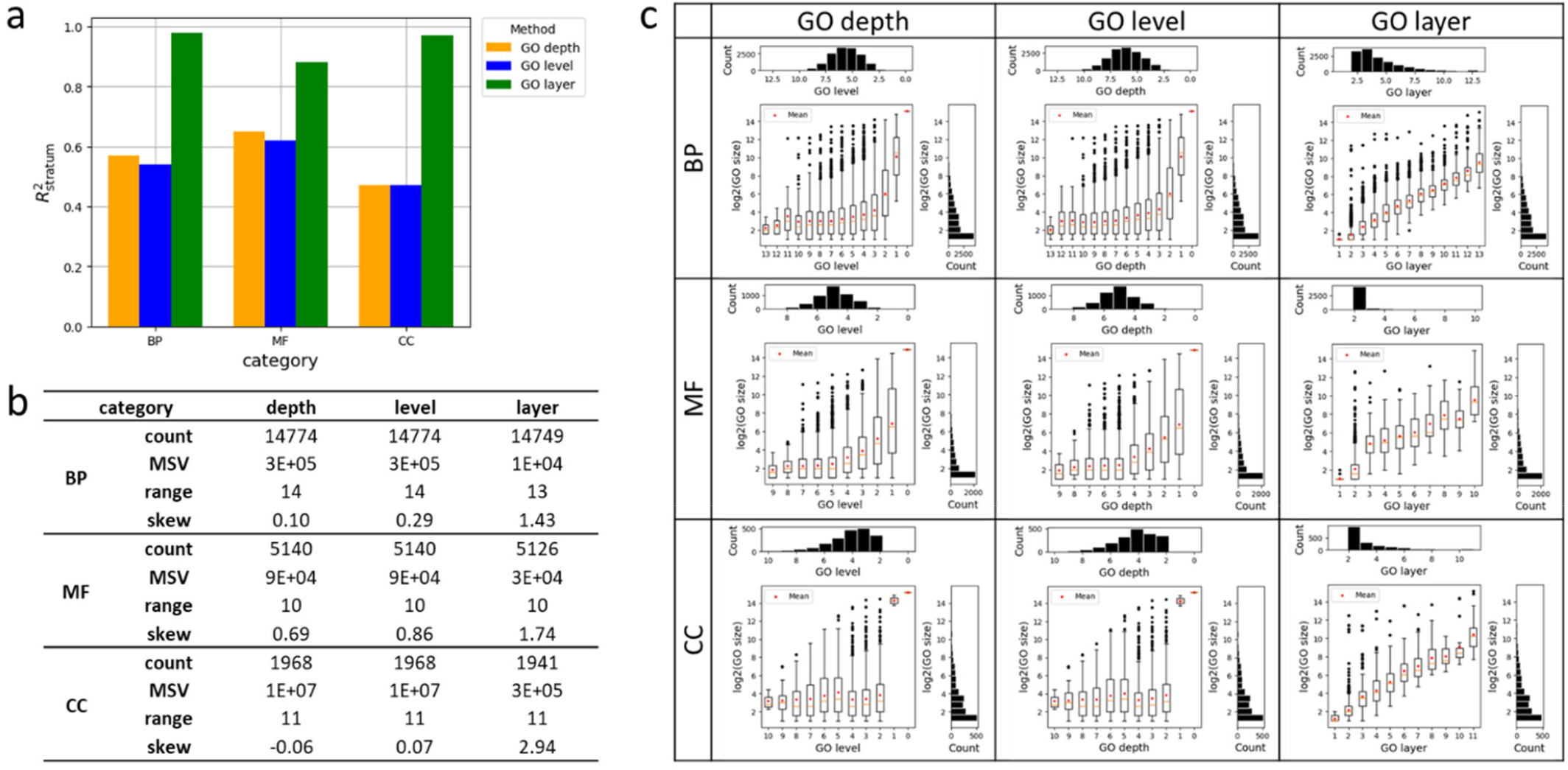
The GOA degree’s R-squared (R-sq.), Aggregation Index (AI) maximization by the optimized Cumulative Density Function (CDF) value in (A) merged GOA, (B) Biological Process GOA, (C) Molecular Function GOA, and (D) Cellular Component GOA.

### The GOALS layer aggregates specific GO terms in a bottom-up manner, forming a multiple layered structure with a pyramid-like distribution of GO terms

The GO layer demonstrates a pyramid-like GO term distribution by folding GO terms from relative abstract-content stratum initially reported (a lower GO depths in relatively abstract stratum) to a specific-content stratum with an increasing degree of inequality towards specific-content GO layer (**Fig. 3**). The pyramid-like distribution of GO layers arises from the degree-based organization of GO terms, which are decomposed from scale-free m-type networks (**Fig. S5**). The polarity of GO term distribution, measured by the Degree of Inequality (DI)(24), reflects the extent to which GO terms are disproportionately concentrated in either specific or abstract strata. A positive DI indicates a skew toward more specific layers. Across all three GO categories, the GO layer stratification results in positive DI values, 0.39 for GO-BP, 0.57 for GO-MF, and 0.46 for GO-CC (**Fig. 3a** and **Table S3**), suggesting a tendency for GO terms to cluster within strata representing more specific biological concepts. Despite this aggregation, the relative specificity of GO terms is preserved, as evidenced by the Positive Pointwise Mutual Information (PPMI)(25) heatmap. Furthermore, Pearson correlation coefficients from the comparison of GO strata show that the GO layer maintains overall granularity, yielding consistently positive values (**Fig. 3b**). The Lorenz curve (**Fig. 3c**) further supports this finding, indicating a positive enrichment of terms toward specific layers. Additionally, the PPMI heatmap demonstrates how granularity is retained by capturing semantic continuity, aggregating GO terms from relatively more abstract positions adjacent to the original strata.

**Figure 3.**
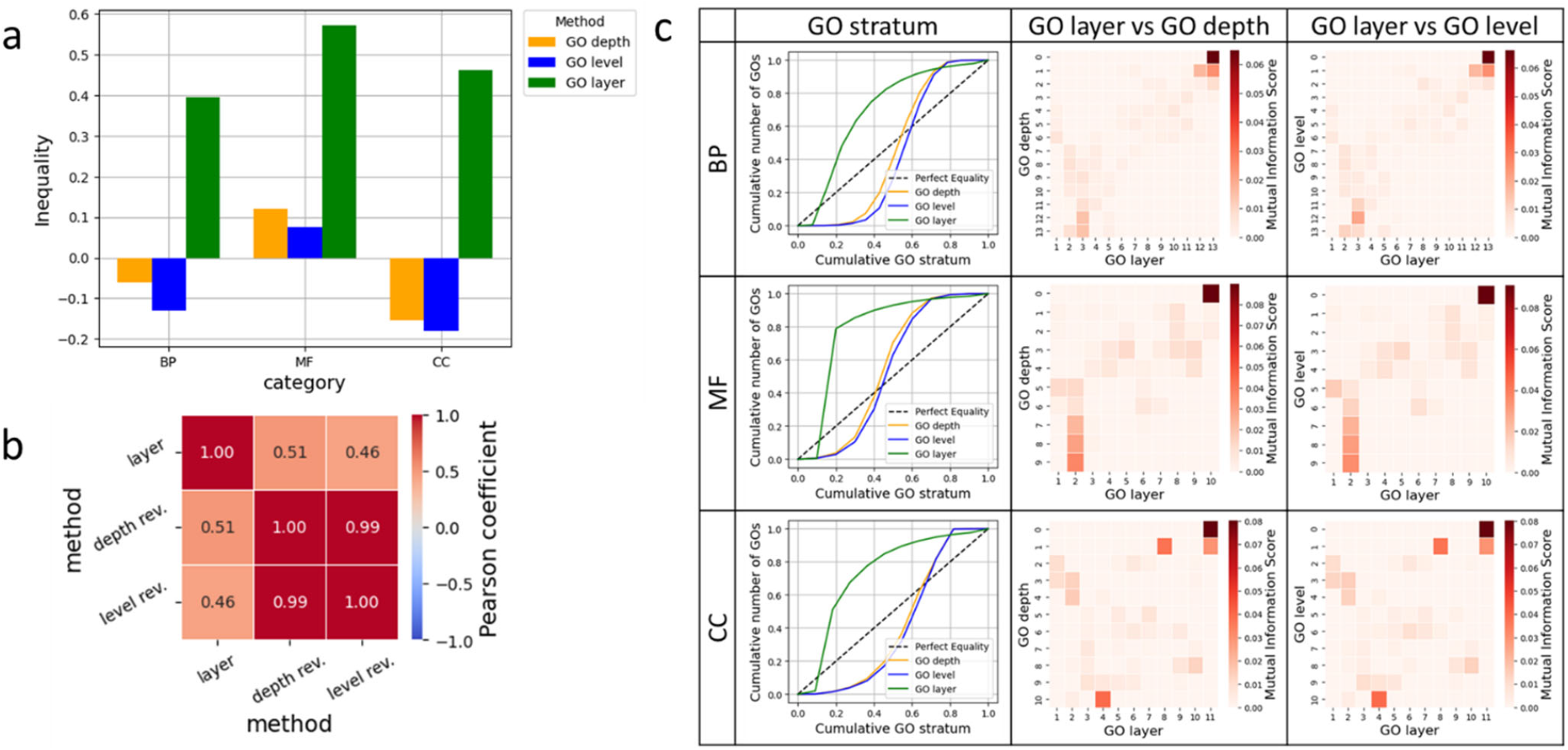
The GOALS layer inequality and its comparison with GO depth and level. (a) The inequality index comparison among GO depth, GO level and GO. (b) The correlations among the three GO stratification methods, GO depth, GO level and GO layer. (c) The Lorenz curve and heatmap depict the similarity comparison of GO layer, GO depth, and GO level using mutual information across GO-BP, GO-MF, and GO-CC.

### The GO map with clustered GO groups represents their semantic subdomains in functional annotation

The GO maps generated from the m-type networks offer an intuitive visualization of the relative similarities among GO terms, based on node2vec embeddings(26) followed by UMAP dimensionality reduction(27) (**Fig. 4**). Clustering quality is assessed using the Silhouette Coefficient (SC)(28), Davies-Bouldin Index (DBI)(29), and Calinski-Harabasz Index (CHI)(30) (**Table S4** and **Fig. S6**).

**Figure 4.**
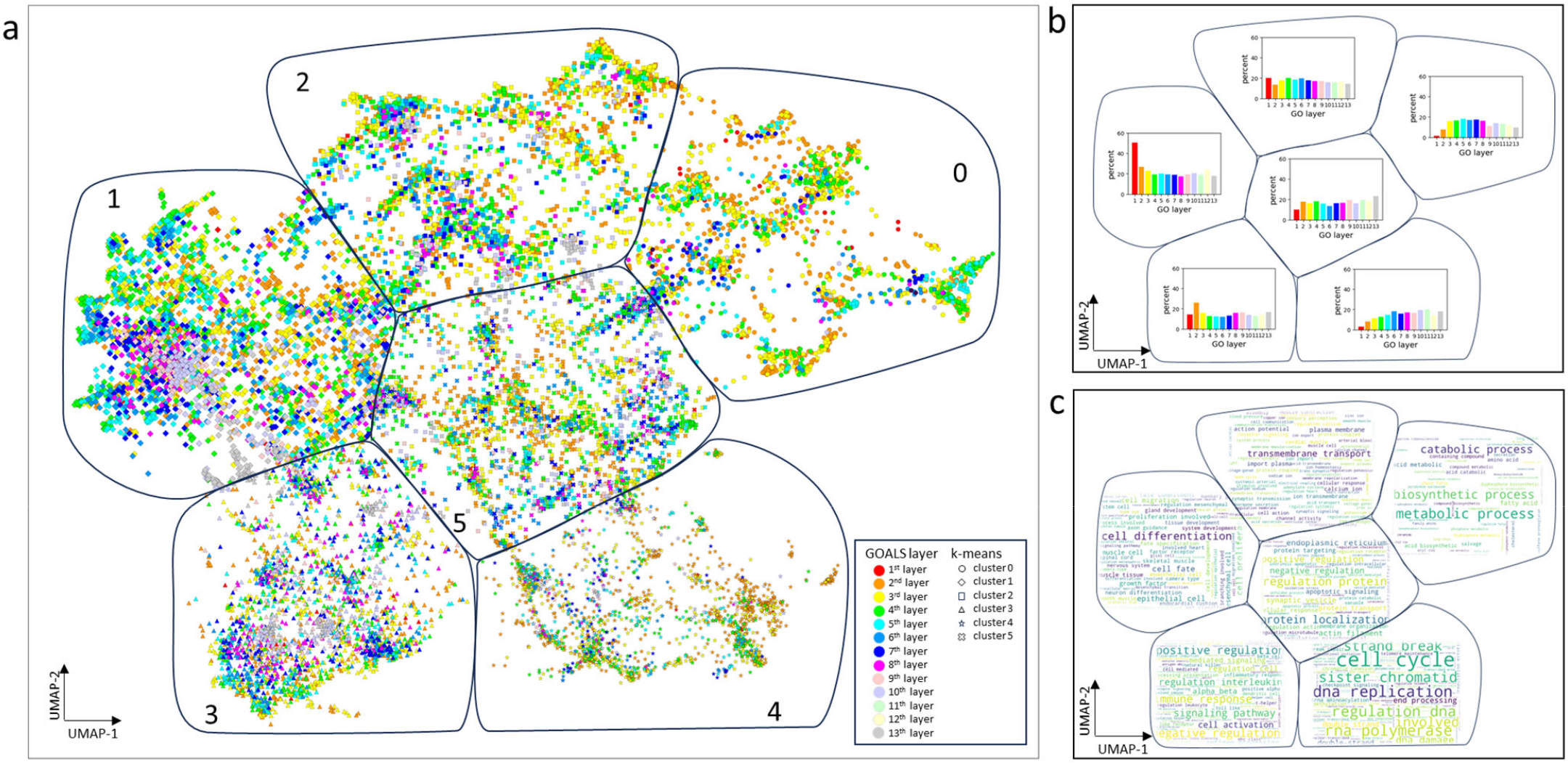
The latent UMAP embeddings of the GO-BP terms of the m-type network. (a) The UMAP projection of the GO terms in the m- type network. (b) Proportions of GO layers within functional groups derived from the clustering algorithms. (c) Word cloud representation of the key terms associated with each identified cluster.

For the GO-BP category, among the four clustering algorithms evaluated, k-means outperforms k-medoids, hierarchical clustering, and spectral clustering. Both the SC and DBI achieve optimal values when the number of clusters is set to 6 using the k-means algorithm (**Table S4**). The UMAP projection, based on 14,749 GO terms, reveals six well-separated functional clusters (**Fig. 4a**). GO-BP terms from different GO layers are broadly and evenly distributed across these clusters (**Fig. 4b**). In the interspersion analysis, GO layer consistently outperforms GO depth and GO level, exhibiting higher interspersion values across the full range of radii from 0.1 to 0.9 (**Fig. S7**). Using the PPMI matrix, distinct biological themes are identified within each cluster (**Fig. 4c**). Cluster #5, centrally positioned in the UMAP space, is enriched for GO-BP terms related to protein regulation and protein localization. Surrounding Cluster #5 are five peripheral clusters: Cluster #0 (metabolic and biosynthetic processes), Cluster #1 (cell differentiation), Cluster #2 (transmembrane transport), Cluster #3 (signaling pathways), and Cluster #4 (cell cycle). The central placement of Cluster #5 suggests that regulatory proteins play a pivotal role in connecting or influencing the biological processes represented in the surrounding clusters, highlighting its importance as a functional hub within the GO-BP landscape.

Similarly, for the GO-MF category, k-means outperforms k-medoids, hierarchical clustering, and spectral clustering among the four clustering algorithms evaluated. Both the SC and CHI achieve optimal values when the number of clusters is set to 6 using the k-means algorithm (**Figure S6** and **Table S4**). The UMAP projection includes 5,126 GO-MF terms and reveals six distinct functional clusters (**Fig. S8a**). GO-MF terms from different GO layers are broadly and evenly distributed across these clusters (**Fig. S8b**). Using the PPMI matrix, distinct biological clusters with their subdomains are identified within each cluster (**Fig. S8c**). Cluster #5, centrally positioned in the UMAP space, is enriched for GO-MF terms related to ligase activity. Surrounding Cluster #5 are five peripheral clusters: Cluster #0 (dehydrogenase and oxidoreductase activities), Cluster #1 (receptor binding and activity), Cluster #2 (transporter activity), Cluster #3 (methyltransferase activity), and Cluster #4 (phosphatase and kinase activity). The central placement of Cluster #5 suggests that dehydrogenase and oxidoreductase activities play a pivotal role in regulating or being regulated by multiple molecular functions represented in the surrounding clusters, highlighting its importance as a functional hub within the GO-MF landscape.

For the GO-CC category, among the four clustering algorithms evaluated, no consistently winning algorithm among k-means, k-medoids, hierarchical clustering, and spectral clustering. The SC achieves optimal values when the number of clusters is set to 2 using the k-means algorithm (**Figure S9** and **Table S4**). The UMAP projection includes 1,941 GO-CC terms and reveals two distinct functional clusters (**Fig. S9a**). GO-CC terms from different GO layers are broadly and evenly distributed across these clusters (**Fig. S9b**). Using the PPMI matrix, distinct biological clusters with their subdomains are identified within each cluster (**Fig. S9c**). Cluster #0 is primarily associated with receptor complexes, while Cluster #1 is enriched for mitochondrial components. Together, these two clusters highlight a fundamental functional partitioning within the GO-CC landscape.

### The GO map with clustered GO functional subdomains in a case study of natural killer cells

Although natural killer (NK) cells comprise a relatively small proportion of the immune cell population, they play a pivotal role in the early immune response by mounting rapid and immediate cytotoxic activity(31). This type of response is particularly critical in the context of leukemia, offering valuable therapeutic potential through engineered modifications of NK cells, such as CAR-NK cell therapy(32, 33). NK cell maturation and differentiation directly impact CAR-NK cell efficacy, safety, and durability(34, 35). Understanding and controlling these processes enables the development of more effective and targeted immunotherapies for cancers and possibly even viral infections(36). The single-cell sequencing data from mouse bone marrow, blood, and spleen tissues provide an opportunity to identify the mechanisms in NK cell maturation and differentiation(19).

The GO map enables intuitive navigation through functional subdomains by incorporating GO layer information, allowing a hierarchical drill-down from abstract concepts to trajectory-specific GO-BP terms related to mature natural killer (mNK) cells. These GO terms are identified based on differentially expressed genes (DEGs) along defined cell developmental trajectories and are further refined through clustering analysis (**Fig. 5**).

**Figure 5.**
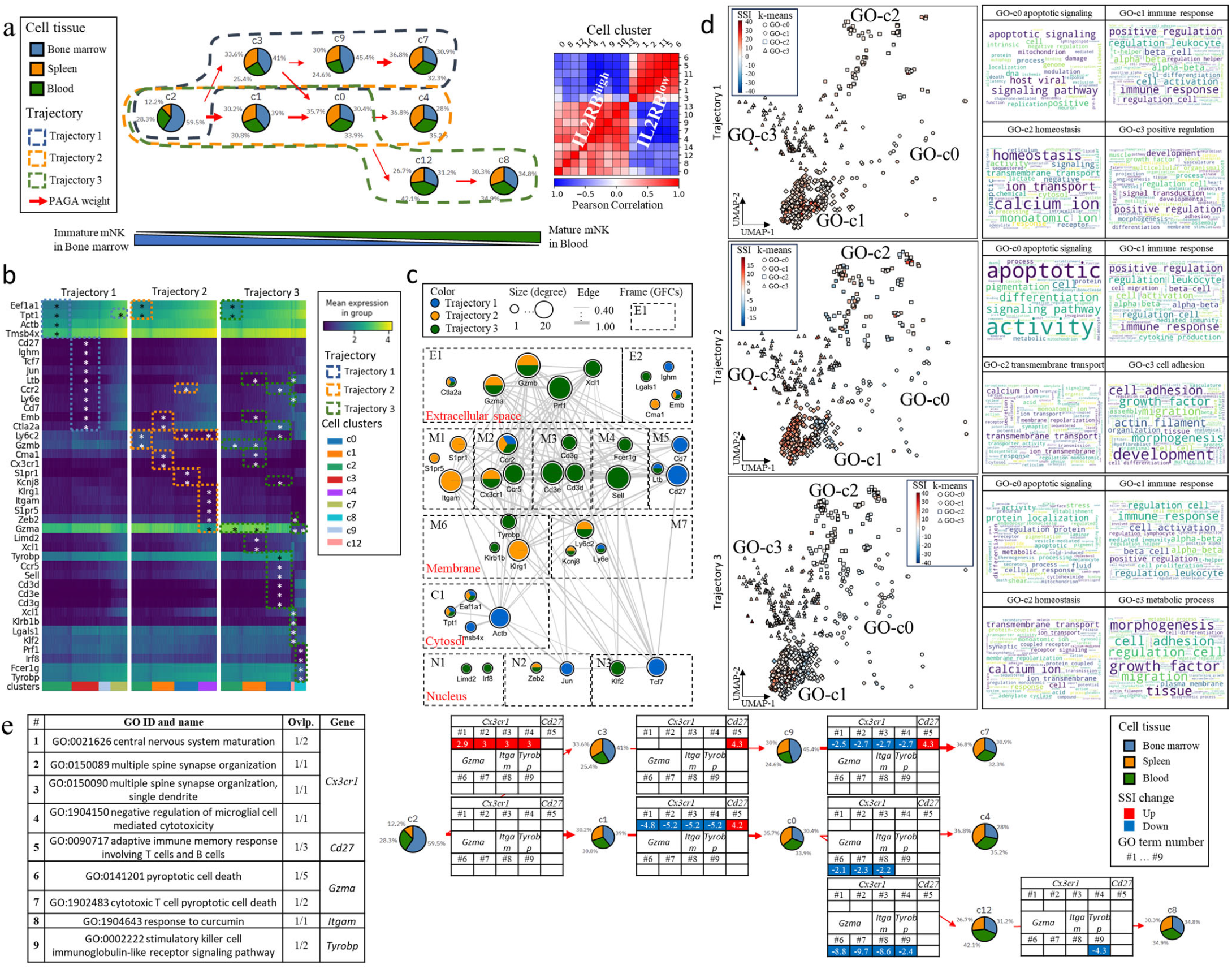
The three major trajectories of mouse natural killer (mNK) cell clusters in a multifurcation model. (a) Two major constituent cell types consisting of 9 cell clusters with different proportions of tissues identified through unsupervised analysis of single-cell transcriptomes. PAGA represents partition-based graph abstraction. (b) Cell-cluster-specific genes across the trajectories identified using differential expression analysis. (c) The inferred network-based trajectory-specific gene signal transduction with gene functional compartments (GFC). (d) The visualization of GO map signal strength index (SSI) and identified GO functional groups. (e) The signal map details of GO-BP map layer 2 GO terms from immune response cluster (GO-c1).

mNK cells are clustered into nine distinct cell populations, forming three cell trajectories. Using biomarker expression (e.g., *IL2RB*) and mNK cell composition across three tissues, bone marrow, spleen, and blood, the directionality of cell maturation from precursor to mature mNK cells is inferred using a PAGA-weighted trajectory map (**Fig. 5a**). DEGs are extracted across transitions between cell clusters, and statistically significant DEGs are visualized in a heatmap (**Fig. 5b**).

A protein-protein interaction (PPI) network is constructed to visualize the relationships among 39 genes identified across the three trajectories. Of these, 31 genes (80%) formed a connected network, demonstrating a high degree of functional interaction (**Fig. 5c**). Within this set, 13 genes are shared across at least two trajectories, while 26 genes are trajectory specific. Subcellular localizations for these genes are inferred using GO-Cellular Component (GO-CC) annotations. The network is further decomposed into 13 Gene Functional Compartments (GFCs) based on functional classifications obtained from GeneCards(37) and WikiGene(38), as introduced in our previous work.

Compared to the GFC-based annotation, the GO map provides a scalable and unbiased GO screening that integrates layered GO panels, enabling visualization of enriched GO functions with their corresponding gene sets. Enrichment analysis results are presented in a global view, allowing users to explore functional clusters and their subdomains (**Fig. 5d**). For example, GO Cluster 1 (GO-c1), which is associated with immune response, emerged as a highly relevant cluster to NK cell differentiation and maturation. By drilling down to a specific GO layer, such as GO-BP layer 2, we identified 9 enriched GO-BP terms, along with their associated gene members.

To provide functional insights along the developmental trajectories, we calculated the Signal Strength Index (SSI) for each GO term, offering a GO-level signal trajectory map aligned with cell cluster transitions (**Fig. 5e** and **Table S5-S7**). In the early divergence between Trajectory 1 and Trajectory 2, *Cx3cr1* is uniquely upregulated during the transition from cell cluster 2 to cluster 3 in Trajectory 1 (**Fig. 5e** and **Fig. S10**). This gene is implicated in central nervous system maturation, synapse organization, and negative regulation of microglial cell-mediated cytotoxicity. Conversely, in the later divergence between Trajectory 2 and Trajectory 3, *Tyrobp* is uniquely downregulated during the transition from cluster 0 to cluster 12 in Trajectory 3 (**Fig. 5e** and **Fig. S10**), suggesting the activity changes in killer-cell immunoglobulin-like receptors (KIRs) modulated by stimulatory killer cell immunoglobulin-like receptor signaling pathway.

All the five genes (*Cx3cr1, Cd27, Gzma, Itgam*, and *Tyrobp* genes) screened by the GO-BP map layer 2 GO terms from immune response cluster (GO-c1) are critical in NK cell maturation, function, and immune regulation supported by existing literature. *Cx3cr1* is associated with late-stage, highly cytotoxic NK cells(39). Loss of *Cx3cr1* impairs NK cell responses to infection and inflammatory signals. *Cd27* marks early to intermediate stages of NK cell maturation(40). Loss of *Cd27* leads to reduced NK cell survival and function, impairing immune responses(41). Mature NK cells shift from *Gzma*^*high*^ to *Gzmb*^*high*^, increasing their direct cytotoxic capacity(42, 43). *Itgam* is a hallmark of terminally differentiated NK cells(44, 45). The decrease of *Itgam* leads to a homeostatic expansion stage of NK cells(45). *Tyrobp* is critical with a dual role in NK cell activation and inhibition(46-48). *Tyrobp* stimulation will enhance cytotoxic signaling in response to MHC target cells. In summary, GOALS effectively identified biologically relevant GO terms in the scRNA-seq of mNK cells.

## Discussion

The GOALS framework offers a novel perspective on Gene Ontology (GO) analysis by introducing a hierarchical GO-layer structure to address longstanding limitations in granularity and interpretability. Unlike conventional GO term fusion, such as ClueGO(6), relying on the kappa index to merge and identify overrepresented GO terms from a set of GO terms, GOALS enables multi-layered structural screening across the GO space. This overcomes the challenge of inconsistent resolution within and across GO strata with significantly reducing the gene size variance within GO term strata, allowing for the systematic identification of biologically meaningful patterns that are often lost in traditional flat enrichment analyses.

A key innovation in GOALS lies in its m-type network architecture, which captures a global topological view of GO terms and their relationships rather than focusing solely on predefined depth, GO level, or k-shell decomposition. This design allows for the detection of hierarchical relationships and functional modules with consistent gene memberships. Importantly, the introduction of an α-percentile thresholding strategy reveals a power-law distribution in GO term connectivity, supporting a pyramid-like organization of GO space. This structure enables GO terms to be aggregated into layers with coherent biological specificity, offering a principled alternative to arbitrary cutoffs.

GOALS not only improves interpretability but also ensures that all enriched terms are preserved and organized by their structural relevance, minimizing the risk of excluding significant terms due to stringent cutoffs, as is common in tools such as DAVID(49), enrichR(50), WebGestaltR(51), and g:Profiler(22). Moreover, the GOALS pipeline can be seamlessly integrated into existing functional genomics workflows, supporting downstream applications, including gene set enrichment, pathway analysis, and disease signature interpretation.

Beyond structural enrichment, GOALS introduces GO map, a latent space by leveraging embedding-derived functional clusters. These clusters capture nuanced relationships among GO terms, informed by co-occurrence patterns in gene annotations. The framework is designed to be flexible, allowing additional dimensions such as GO metadata, expression specificity, or evidence codes to refine network layouts. Furthermore, domain-specific networks, such as those defined by protein-protein interactions, transcriptional regulation, or shared drug targets, can be overlaid to enhance biological relevance.

GOALS opens new avenues for condition-specific GO term comparison, facilitating the distinction between triggering GO terms (those likely initiating biological responses) and passenger GO terms (those resulting from downstream effects). This causal decomposition may benefit from the integration of transcription factor binding data, genetic variant mapping, and other regulatory annotations, enabling a more mechanistic understanding of enriched GO signals.

The GOALS framework has strong potential for multi-omics integration, wherein network fusion could elucidate global activity patterns and causal pathways across different biological layers. The ability to infer structured, interpretable, and biologically grounded enrichment patterns positions GOALS as a powerful tool for next-generation systems biology and precision medicine applications.

## Methods

### An overview design of the GOALS framework

We present the GOALS framework, a four-stage data integration pipeline (**Fig. S1**) designed to generate and prioritize layered Gene Ontology (GO) terms for enhanced functional genomics analysis. This framework emphasizes interpretability, scalability, and biological relevance throughout each integration stage. In stage 1 GO layer decomposition using the latest GO release (2025/02/06), GO terms are first processed through the GO Consortium’s database. A co-membership (m-type) GO-to-GO network is constructed and optimized, with quality control measures ensuring conformity to a power-law distribution via a linear regression test. This step forms the basis for a stratified, layered GO term structure. In stage 2 the GO map generation using embeddings, the GO depth and level are used as the baseline to compare the GO layer performance based on the GO size’s power law distribution fit in a linear regression and inequality test. The GO embeddings are yielded from the m-type GO-to-GO networks using node2vec. In stage 3 genomics profile overlay, the differentially expressed genes are evaluated based on the logarithm-transformed fold changes, log(FC) and statistical significance comparing different biological conditions. A Signal Strength Index (SSI) is calculated for each GO term by aggregating expression signals of its gene members, thereby prioritizing biologically meaningful GO terms within each layer. In stage 4 comparative phenotype analysis, the sample phenotype (or subset of cell populations in cell trajectory in single-cell data) is integrated to assess the differential significance of SSI across conditions. Layer-specific GO terms are then validated using literature-supported gene’s function to disease associations to ensure interpretive robustness.

### Extract the GO information and “is-a” relationship and construct the m-type GO-to-GO network

The most up-to-date GO was downloaded from the GO consortium (https://www.geneontology.org/). We processed GO term information, including the GO identifier, name, description, category, organism, link and “is-a” relationship, from the “.obo” file. Gene-specific GO annotations were extracted from the “.gal” file. According the “is-a” relationship, all the child annotated genes was aggregated to their parent GO terms in a recursive loop to ensure the parent node fully cover the genes belonging to their child nodes. To assess the confidence of each m-type GO-to-GO pair, we applied the probability mass function (PMF) of hypergeometric distribution as published in PAGER(17). Only over-represented relationships were considered, with under-represented associations assigned a value of zero. The PMF is defined in Equation 1.

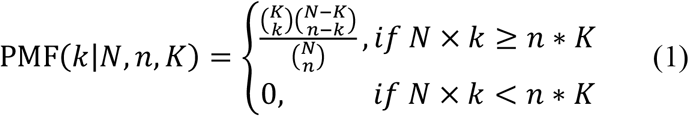

Where *N* was the total number of genes covered in GO terms, *n* was the number of genes annotated to the GO term GO_*i*_, K was the number of genes annotated to GO term GO_*j*_, and k was the number of overlapping genes between GO_*i*_ and GO_*j*_.

### Perform quality control of m-type networks maximizing the product of the power-law distribution fit and the Aggregation Index of m-type networks

To assess the fit of the power law distribution for the GO gene numbers, we used the coefficient of determination from a linear regression of the logarithm transformed GO degrees in the network and logarithm transformed GO frequencies, denoted as 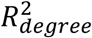. To evaluate the quality of network coverage, the Aggregation Index (AI) (52) was employed based on the GO term’s coverage in the largest network. Both the 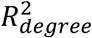 value and AI ranged from 0 to 1, with higher values indicating better network quality. To balance network quality by maximizing the R^2^ value while maintaining adequate coverage, we introduced the AI-adjusted R^2^ Index (ARI) in Equation 2.

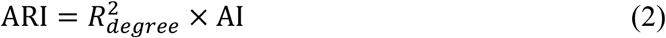

We then screened the optimal cutoff for the -log(PMF) values within a range of 0 to 10, optimizing both the network’s fit to the power law and its coverage.

### Stratify GO layers using a scalable threshold to determine the set of extended GO terms

In the Assignment and Expansion steps, a new layer was assigned by incrementing the current layer by 1, extending to a set of GO term candidates, denoted by *N*. These terms had degrees equal to or below a predefined threshold, determined by the *α*-percentile of the degree distribution among candidate GO terms, as specified in Equation 3:

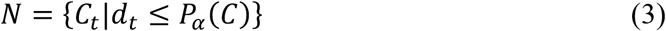

Where *d*_*t*_ represented the degree of the GO term indexed by *t*, and *P*_*α*_*(C)* denoted the *α*-th percentile of degrees among all the non-interactive GO term candidates in the pool *C*. The parameter *α* is the scale factor varied from 0 to 1, controlling the threshold from stringent to relaxed inclusion criteria.

### Optimize GO layer stratification by maximizing the product of the power-law distribution fit and a baseline-match-adjusted index based on layer range

To optimize the GO layers stratification using the α percentile cutoff, we introduced the baseline-match- adjusted *R*^2^ index (BRI). The *R*^2^ ensured power law distribution and penalized deviations when the GO layer exceeded or fell below the established GO stratum baseline. BRI incorporated both and a baseline match score for evaluating the GO layer range. To assess the fit of the power law distribution for the GO gene numbers, we used the coefficient of determination 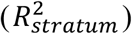 from a linear regression of the logarithm transformed GO frequency in the network and total number of distinct GO layers. All the GO terms (with a total of *T*) were assigned with layer number in a list, denoted as *ℒ = [layer*_1_, *layer*_2_,…, *layer*_*T*_], where *t* is an index of GO term. The total number of distinct GO layers was defined in Equation 4.

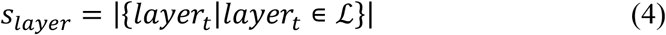

The GO stratum baseline (*s*_*baseline*_) was defined as the average of the total number of distinct strata from both GO depth and GO level in Equation 5.

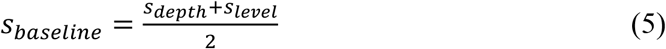

Where the *s*_*depth*_ represented the total number of distinct GO depth in the GO list with depth number, *𝒟= [depth*_1_, *depth*_2_,…, *depth*_*T*_] in Equation 6.

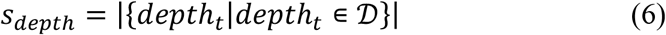

Similarly, the *s*_*level*_ represented the total number of distinct GO levels in the GO list with level number, *θ = {level*_1_, *depth*_2_,…, *depth*_*T*_*}* in Equation 7.

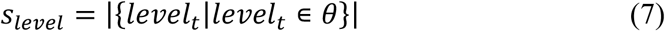

To compare the number of GO layers with the stratum baseline, the baseline-match-adjusted *R*^2^ index (BRI) was computed to evaluate the performance of the GO layer stratification in Equation 8.

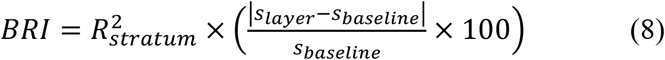

We will select the optimized *α* percentile cutoff maximizing BPI.

To evaluate reduction of GO term’s gene size variance within the same GO layer, we applied the measurement, the mean of sample variance (MSV) across distinct layers, as shown in Equation 9.

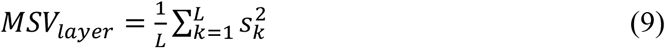

Where *L* was the total number of GO layers and 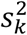 represented the sample variance of gene sizes among GO terms within the *k*-th layer defined in Equation 10.

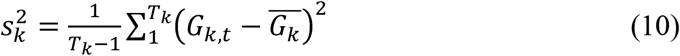

Where *T*_*k*_ was the total number of GO terms within the *k*-th layer. *G*_*k*,*t*_ represented the gene size of GO term indexed by *t* among GO terms within the *k*-th layer defined in equation 11. 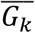 represented the average number of genes of GO terms within the *k*-th layer defined in Equation 11.

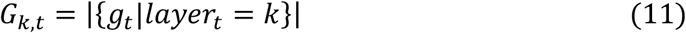

Where *g*_*t*_ represented genes belonging to a GO term indexed by *t* and *layer*_*t*_ *= k* indicated that the corresponding GO term belonged to the *k*-th GO layer.

### Perform inequality test for GO distribution polarity in GO strata

To evaluate the GO terms distribution polarity in the GO layer strata, we performed the inequality test using the Lorenz curve and an inequality index. To order GO stratum from specific to abstract, GO depth and level were reverse ordered. The cumulative number GO was calculated based on the GO falling into the GO strata from specific to abstract. The Lorenz curve was generated accordingly to compare the baseline of perfect equality to evaluate the inequality of the number of GO terms enriched towards specific or abstract stratum, ranging from -1 to 1. We employed the Degree of Inequality (DI) of the GO layer in measuring signal inequality, as previously published(24) and defined in Equation 12.

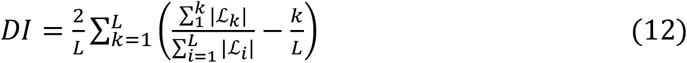

Where *L* was the total number of distinct GO layers. *i* denoted the value of the GO layer ranging from 1 to *L*. |*ℒ*_*k*_| denoted the cardinality of the layer specific to the *k*-th GO layer in Equation 13.

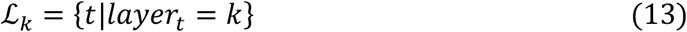

Where *ℒ*_*k*_ denoted the set of indices for those GO layer is *k*.

To enable comparisons among the GO term stratum from different approaches, GO depth, GO level and GO layer, we generated the mutual information-based similarity matrix. The positive pointwise mutual information (PPMI) matrix was calculated based on a similarity matrix of gene members of GO terms between the two types of GO stratum, such as GO layer versus GO depth, denoted as *mtx* ∈ *ℝ*^*LXD*^, where *L* was the total number of distinct GO layers and *D* was the total number of distinct GO depths.

The similarity matrix was generated using a similarity score from the Jaccard index and overlap coefficient to mitigate the GO size bias. The similarity score was calculated in Equation 14.

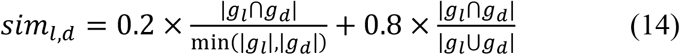

Where *g*_*l*_ and *g*_*d*_ represented genes belonging to a GO term indexed by *t* and *d* respectively. The joint probability was calculated in Equation 15.

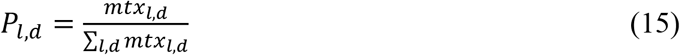

Where *l* was the index of the distinct layer number and *d* was the index of the distinct depth number. The marginal probability of the GO layers in rows in Equation 16.

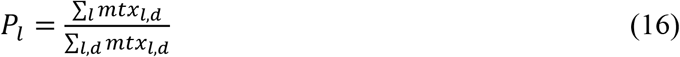

The marginal probability of the GO depths in columns in Equation 17.

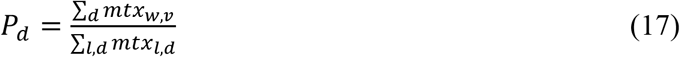

The PPMI for comparing GO layers and GO depths was calculated as defined in Equation 18.

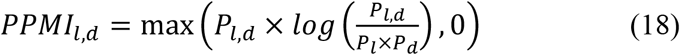

The PPMI matrix was used to generate a heatmap that reveals the similarities and granularity of GO term distributions across different GO stratification methods.

### Generate the embedding of the m-type network and perform dimension reduction

To generate embedding for the network, we used the node2vec algorithm for learning continuous representations of GO terms in the m-type networks. Node2vec captured m-type network structural information by optimizing a neighborhood-based objective that is sensitive to both local and global graph properties. Then, we applied Uniform Manifold Approximation and Projection (UMAP) to transform embeddings into a two-dimensional representation while maintaining the relationships between nodes. The results of UMAP were used to visualize the GO term in two-dimensional space.

To quantitatively assess the distribution and overlap of GO terms within a given stratum (e.g., GO depth, GO level, and GO layer), we computed the interspersion score, which captures how widely and evenly terms were distributed in the UMAP latent space. This metric was based on pairwise proximity and spatial dispersion of GO terms across different GO strata. Given two GO term strata, such two distinct GO layers denoted by *ℒ*_*k*1_ and *ℒ*_*k*2_, and their embedding positions 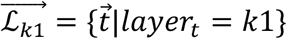 and 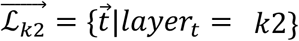 were computed using the UMAP representation of the GO term. The interspersion of the two GO layers *k*1 and GO layer *k*2 given the radius *r*, denoted by *ISP*_(*k*1,*k*2,*r*)_, was calculated in Equation 19.

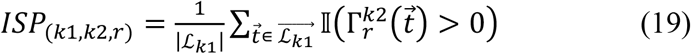

Where 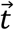 was the point in the set of embedding 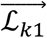 of the GO terms in the GO layer 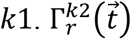 denoted the set of neighbors of the point 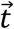within radius *r*, drawn from the set of GO terms in GO layer *k*2. 𝕀(·) was the indicator function, which returned 1 if the condition was true and 0 otherwise.

### Segregate GO and extract keywords using clustering algorithms

To group GO terms into functional clusters within the two-dimensional UMAP latent space, we performed four clustering algorithms for the embeddings, including k-means, k-medoids, hierarchical clustering and spectral clustering. Cluster quality was assessed using the silhouette coefficient (SC)(28), Davies Bouldin Index (DBI)(29), and Calinski-Harabasz index (CHI)(30) to determine both the optimal number of clusters and the best performed algorithms. The number of cluster (k) was set from 2 to 9, and the optimal number of clusters was selected based on the highest SC and lowest DBI, indicating more coherent and well-separated clusters. The GO names within the same cluster were used to calculate the word or phrase relative frequency to reflect represented terms for the cluster. We removed the stop words and punctuations, vectorize the name to extract the bigrams, calculate their cluster-specific frequency for each cluster. If the one word was a substring of extracted bigram that passed the frequency cutoff, this one word was removed. The words from multiple clusters underwent the PPMI analysis. The word frequency matrix was a *w* as the word vectors and a *v* as the GO clusters, denoted as *mtx* ∈ ℝ^*w*×*v*^. The joint probability was in Equation 20.

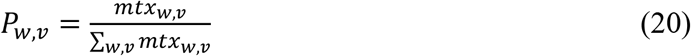

The marginal probability of the words in rows in Equation 21.

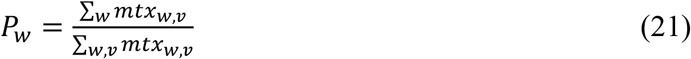

The marginal probability of GO clusters for columns in Equation 22.

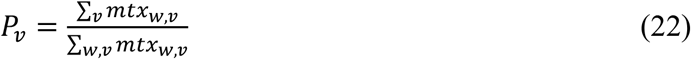

The PPMI was calculated given in Equation 23.

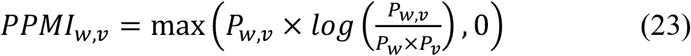

The PPMI matrix was used to generate a word cloud that highlights key concepts, with font size proportional to their significance, using the wordcloud Python library.

### Filter DEGs using fold change and statistical significance

The differentially expressed genes (DEGs) were filtered based on the log(FC) and statistical significance by comparing different biological conditions. In the study of the natural killer (NK) cell, we extracted the three trajectories based on the PAGA in the published work. The DEG list was generated by applying a threshold of absolute log(FC) ⩾ 1 and a Wilcoxon test p-value ⩽ 0.01, along with additional filtering criteria as detailed in our previous work.

### Perform enrichment analysis to generate a list of overrepresented GO terms

Enrichment analysis was performed to identify overrepresented GO terms among DEGs. We utilized the PAGER API to retrieve the list of enriched GO terms. For each GO term, we aggregated the log(FC) values of DEGs overlapped with GO term gene members to compute a GO-level log(FC) values. The significance of enrichment was assessed using the hypergeometric test, and the strength of this significance was expressed as the negative logarithm of the p-value. The signal strength index (SSI) for each GO term was calculated by multiplying the GO-level log(FC) and the negative logarithm transformed p-value, as shown in equation 24.

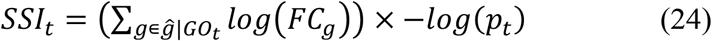

Where *ĝ* |*GO*_*t*_ denoted the set of DEGs that are members of the GO term indexed by *t. logFC*_*g*_ was the log(FC) of the gene *g* as a member of GO term indexed by *t*. −*log(p*_*t*_) denoted the strength of the statistical significance of GO term indexed *t*.

### Evaluate the statistics significance of layer-specific GO terms

To assess the significance of GO’s SSI values across biological conditions, we employed the student’s *t*- test. In the context of scRNA-seq data, enriched GO terms were grouped according to cell cluster-specific expression patterns along with their corresponding SSI values. The *t*-test was then used to identify GO terms exhibiting statistically significant SSI changes across cluster transition, enabling the prioritization of functionally relevant GO terms that potentially drive transitions along multiple cell trajectories.

### Validate layer-specific GO terms and gene members using literature-based evidence

To support the biological relevance of layer-specific GO terms and associated gene members, we conducted literature-based validation using the Biomedical Entity Expansion, Ranking, and Exploration (BEERE) tool(52). Gene symbols, alternative gene names, and GO term’s key words were extracted and used to query the literature for associations with specific biological conditions. BEERE results included disease-relevant publications, along with their corresponding PubMed IDs, providing citation-based evidence to validate the functional importance of the identified GO terms and genes.

### Single-cell collection and sequencing for mouse samples

Single cells were obtained from mouse bone marrow, blood, and spleen tissues. The isolation of mouse natural killer (mNK) cells was carried out using the CITE-seq (Cellular Indexing of Transcriptomes and Epitopes by Sequencing) method, which involved the utilization of antibodies targeting CD11b, CD27, NK 1.1, TCRb, and CD122. This process was performed using three 10x Genomics single-cell lines, each incorporating three distinct hashtags (antibodies) to differentiate the tissue sources.

### Single-cell RNA-seq data processing

The mouse single-cell samples were processed using Cell Ranger v4.0.0, and read alignment was performed with the GRCm38 mouse genome. We conducted filtering to remove low-quality cells and genes based on two criteria: (1) cells with expressed gene (counts larger than 0) less than 200, and (2) expressed genes less than three cells. Since all the samples were processed using the same 10x Genomics platform, we didn’t perform batch effect correction.

### The construction of Protein-Protein Interaction (PPI) network from DEGs and the gene subcellular map

After identifying the DEGs in the selected cluster and comparing them to the reference clusters within the trajectory, we proceeded to retrieve protein-protein interactions from the STRING database (53), employing a confidence score cutoff of 0.40, which represents a moderate confidence level.

### Pseudotime inference and the signal curve plot of GFCs in trajectory

The differentiation trajectory of all mNK cells was inferred using the diffusion map algorithm following the Scanpy workflow (54). We chose the c2 NK cells as the root due to their higher proportion among bone marrow mNK cells. The diffusion pseudotime was calculated using the scanpy.tl.dpt function. To generate the signal curve along pseudotime in each trajectory, we computed the average gene expression for each GFC. Subsequently, we applied a Savitzky–Golay filter (55) in the Scipy library to smooth the signal curve. The window size was set to 601, which roughly corresponds to 25 intervals along the trajectory, and a polynomial order of 3 was employed to capture convolution coefficients in the fitted polynomials.

