## supplementary for "GOALS: Gene Ontology Analysis with Layered Shells for Enhanced Functional Insight and Visualization"

### Figures

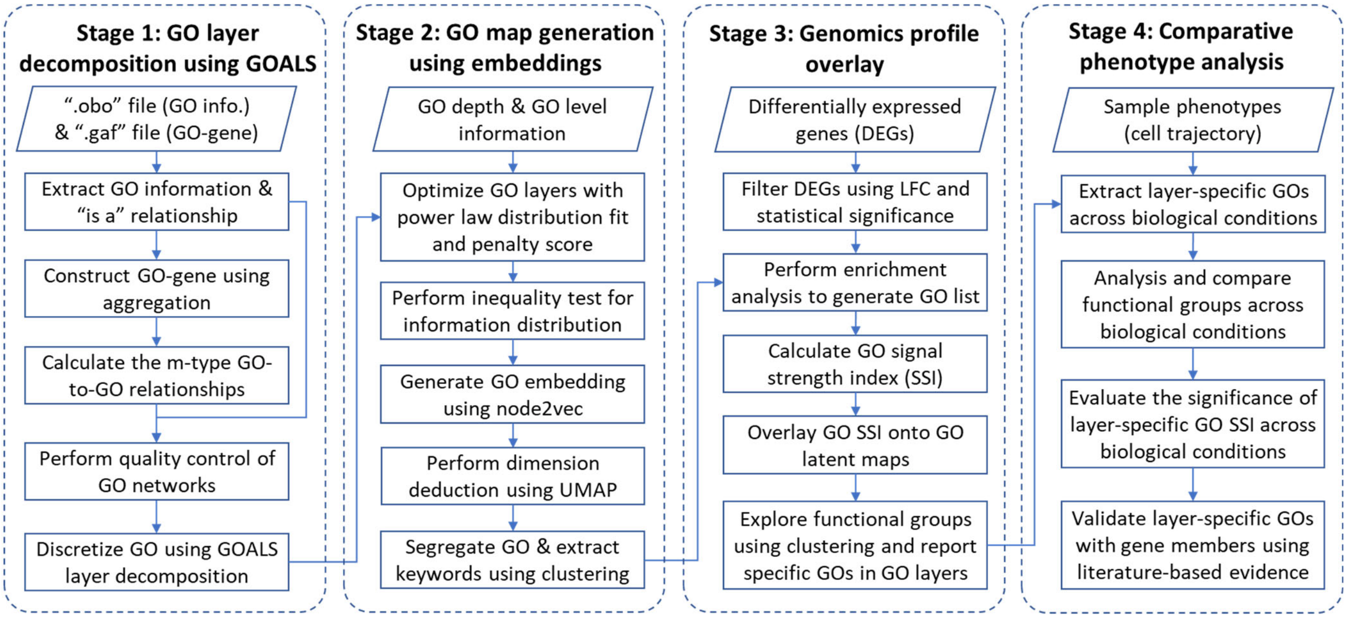


**Figure S 1** The four-stage data integration framework for generating and prioritizing layered GO terms to enhance the interpretation and visualization. The input dataset consists of GO terms from the 2025/02/06 release.

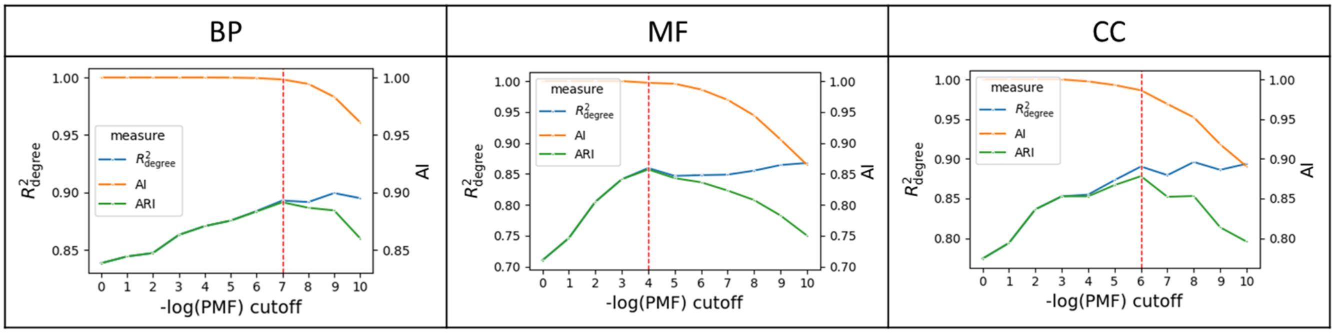


**Figure S 2** The curves of R-squared (R-sq.) of the power-law distribution fit, the Aggregation Index (AI) of networks, and their product ARI across Biological Process (BP), Molecular Function (MF), and Cellular Component (CC) categories. The negative logarithm transformed probability mass function, the cutoff of -nlog(PMF) is indicated by a red dashed line.

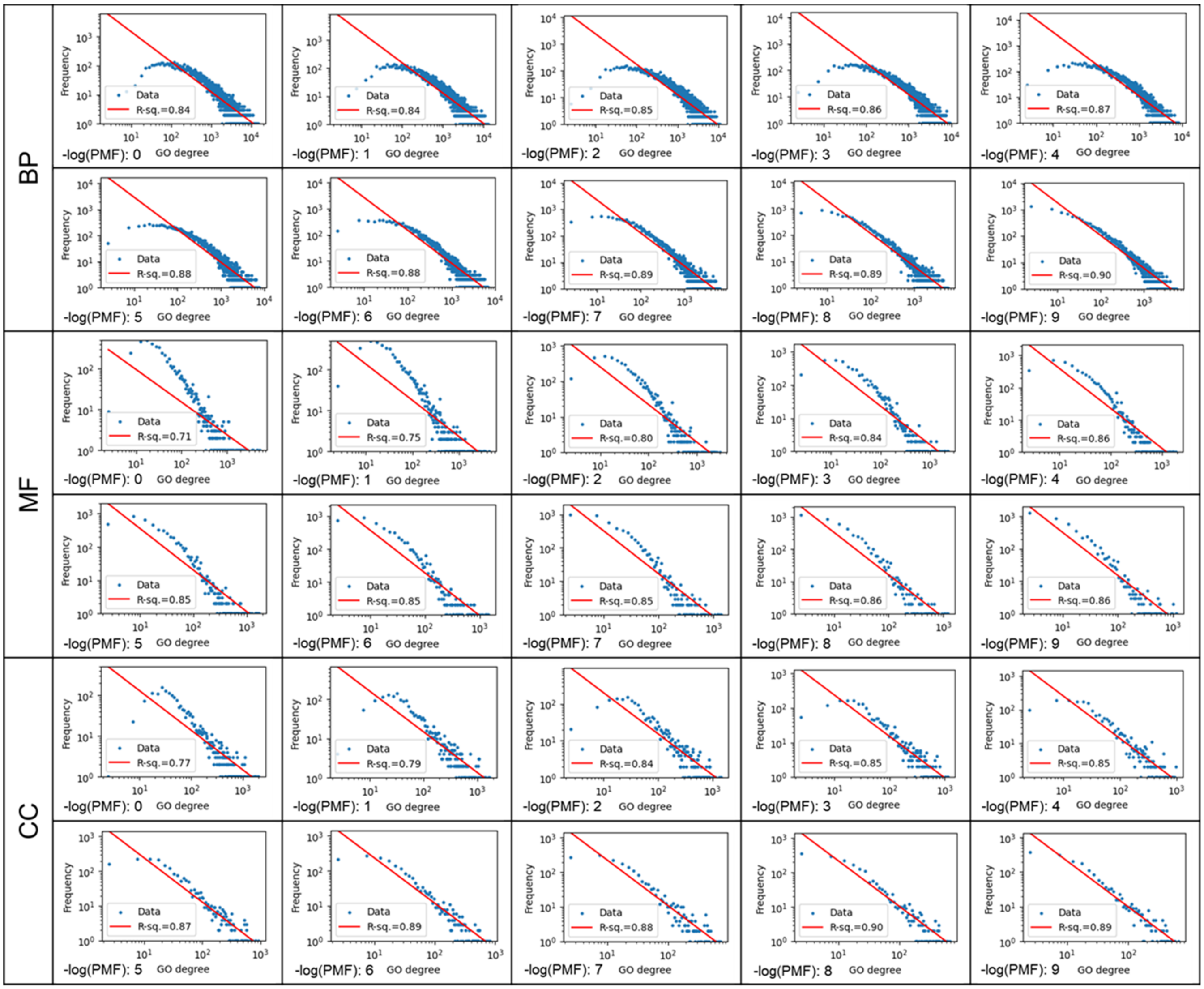


**Figure S 3** The cutoffs of –log(PMF) and the m-type network $R^2_{degree}$ for the power law distribution. The –log(PMF) ranges from 1 to 9 across Biological Process (BP), Molecular Function (MF), and Cellular Component (CC) categories.

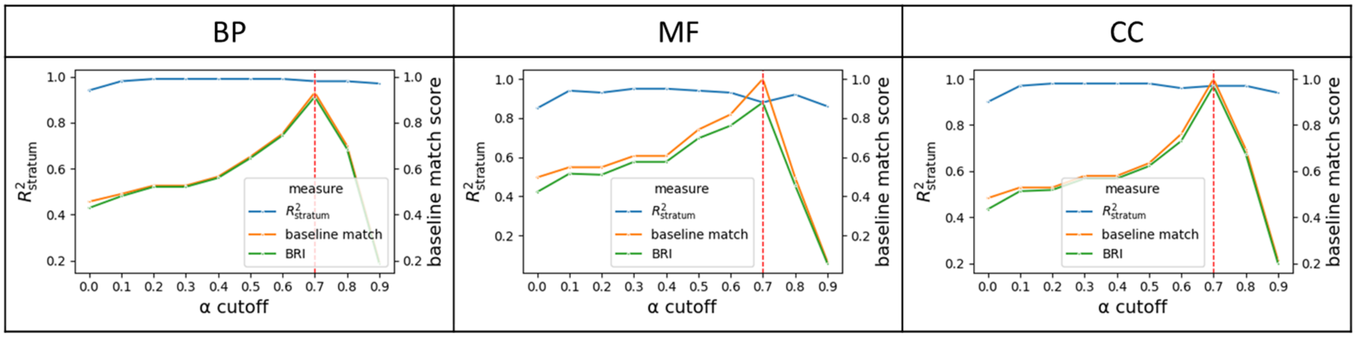


**Figure S 4** The curves of the R-squared ($R^2_{layer}$) of power law distribution fit, the baseline match score from stratum numbers, and their product across GO-BP, GO-MF, and GO-CC categories. The alpha cutoff is indicated by a red dashed line.

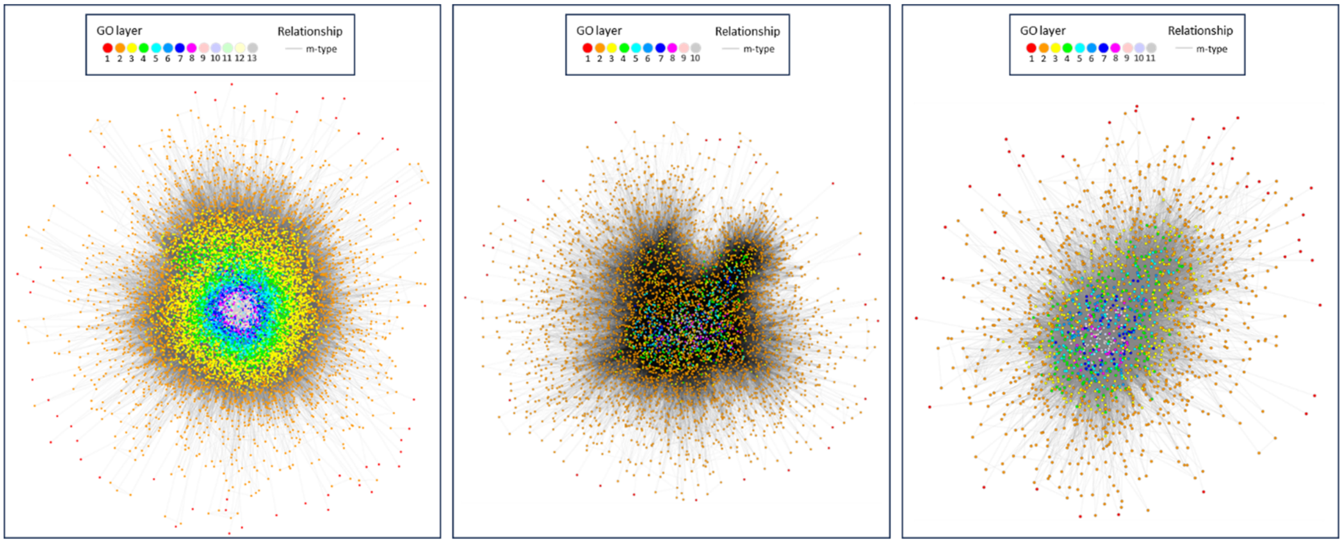


**Figure S 5** The overall view of the GO layer assignment in the m-type networks in the GO categories, including GO-BP, GO-MF, and GO-CC. The color represents the GO layer number. The edge represents the m-type GO-to-GO relationship. The GO-BP network consists of 14,749 GO-BP terms and 2,266,821 m-type GO-to-GO relationships. The GO-MF network consists of 5,126 GO-MF terms and 145,424 m-type GO-to-GO relationships. The GO-CC network consists of 1,941 GO terms and 49,303 m-type GO-to-GO relationships.

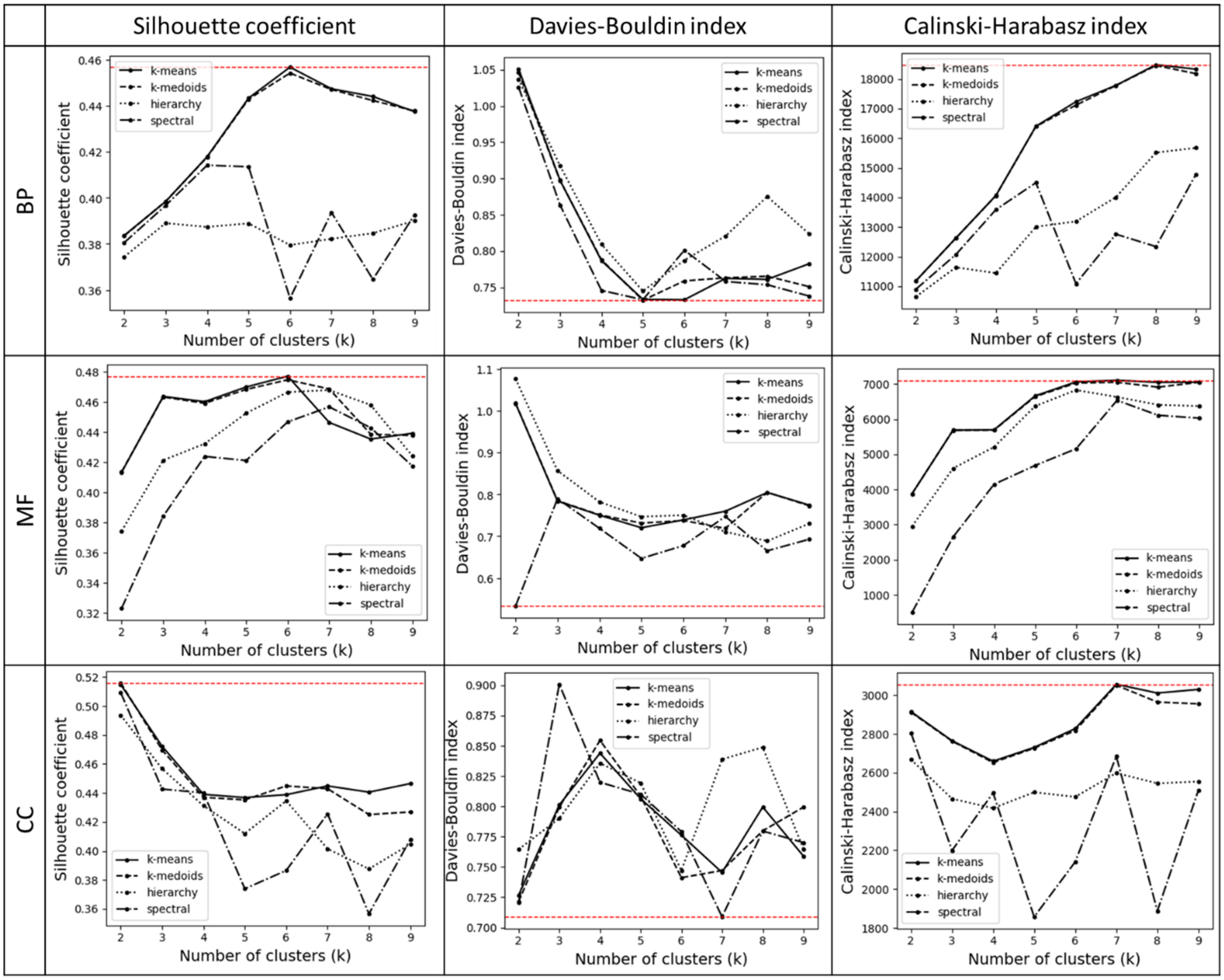


**Figure S 6** The k cluster selection in optimizing the cluster quality using silhouette coefficients (SC), Davies-Bouldin index (DBI), and Calinski-Harabasz index (CHI) in the categories of (a) GO-BP, (b) GO-MF, and (c) GO-CC.

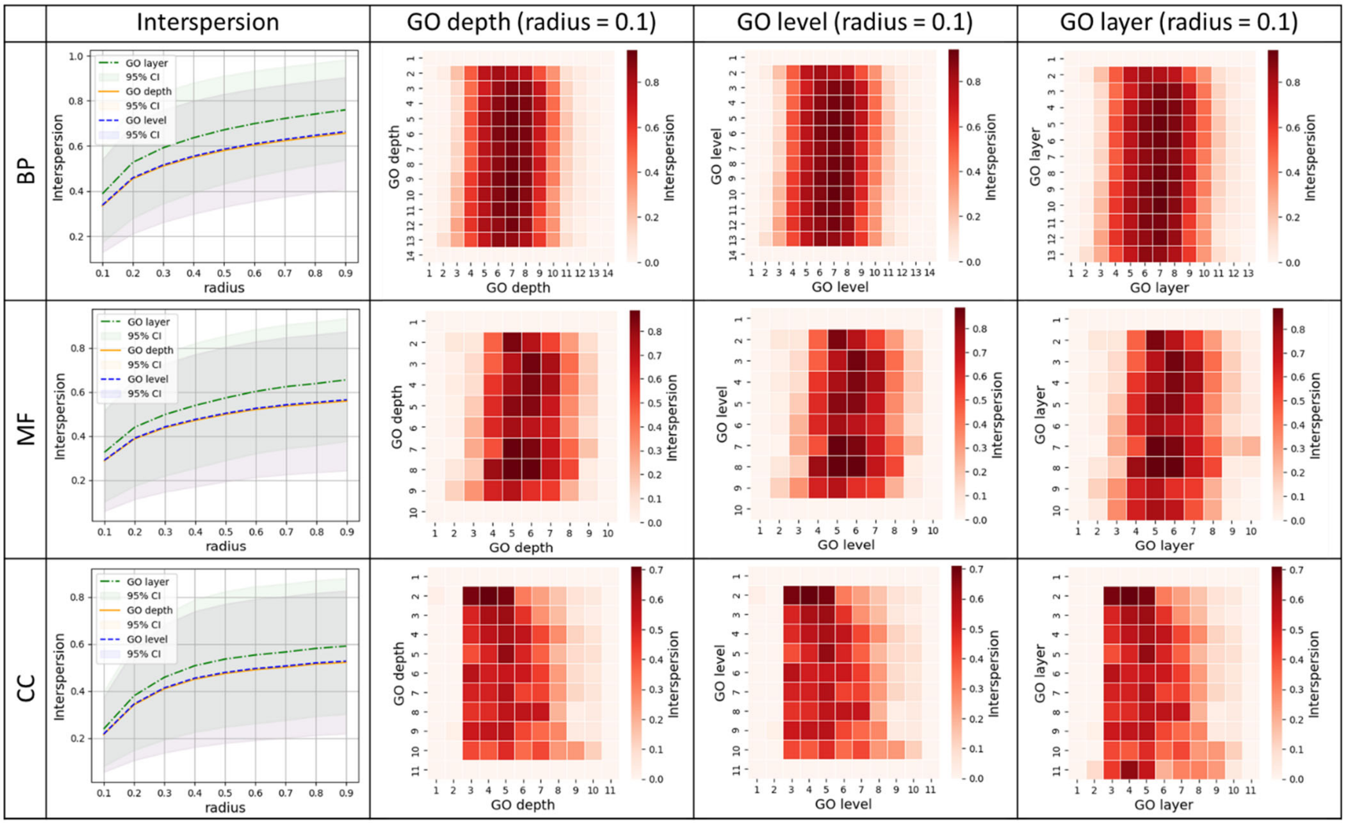


**Figure S 7** Comparison of GO stratum interspersion across three stratification approaches, GO depth, GO level, and GO layer, within the three GO term categories. The first column presents interspersion curves computed over a range of radiuses from 0.1 to 0.8, showing the mean pairwise interspersion values for each GO stratum along with their corresponding 95% confidence intervals. The second to fourth columns display interspersion heatmaps for GO depth, GO level, and GO layer, respectively, providing an example when the radius value is set to be 0.1 across the three GO term categories.

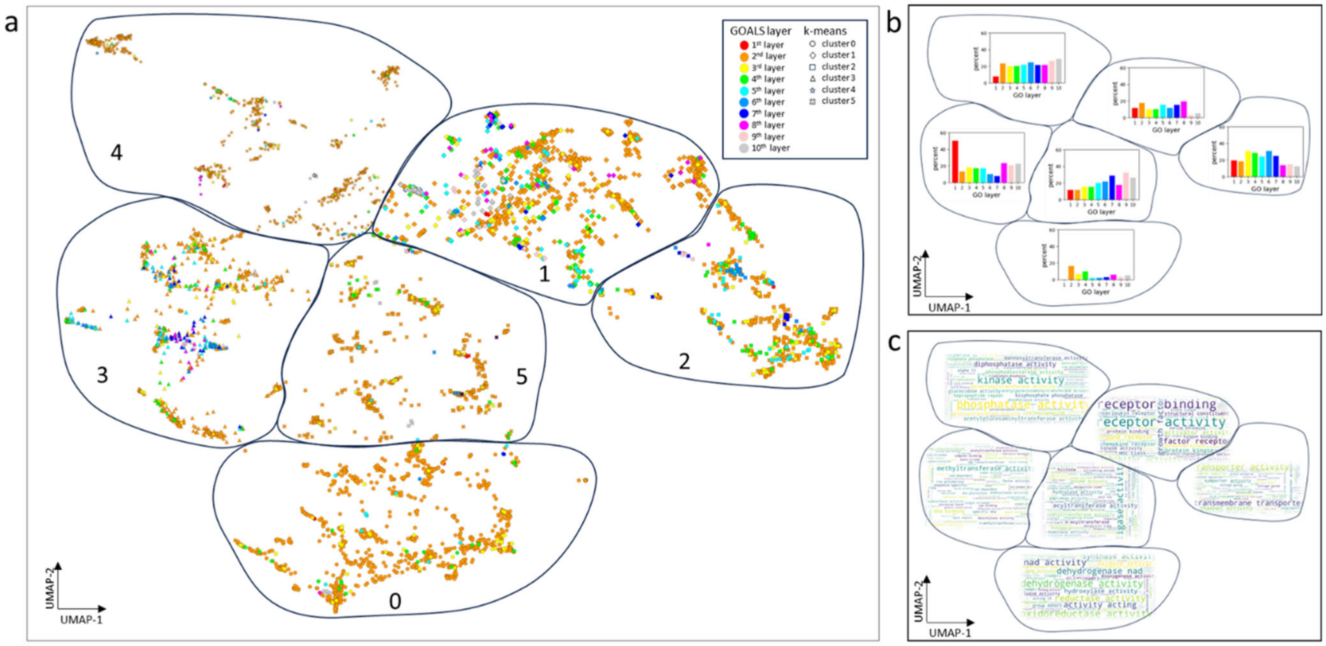

**Figure S 8** The latent UMAP embeddings of the GO-MF terms of the m-type network. (a) The UMAP projection of the GO terms in the m-type network. (b) Proportions of GO layers within functional groups derived from the clustering algorithms. (c) Word cloud representation of the key terms associated with each identified cluster.

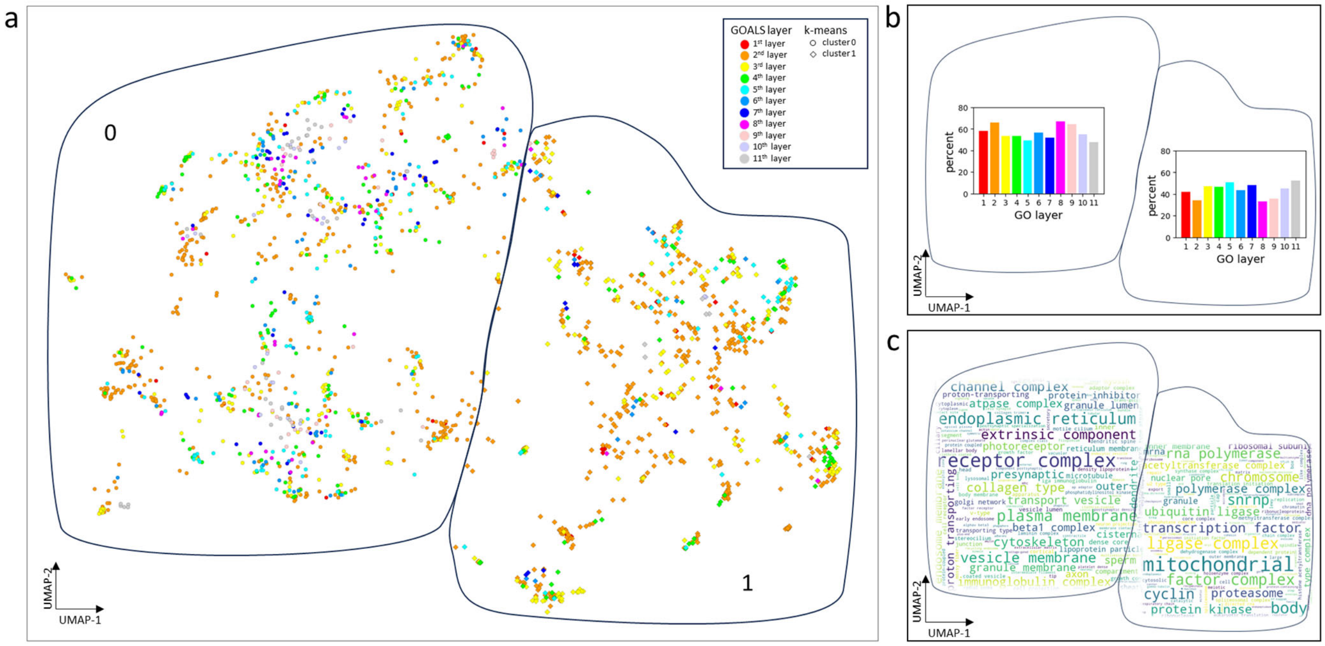


**Figure S 9** The latent UMAP embeddings of the GO-CC terms of the m-type network. (a) The UMAP projection of the GO terms in the m-type network. (b) Proportions of GO layers within functional groups derived from the clustering algorithms. (c) Word cloud representation of the key terms associated with each identified cluster.

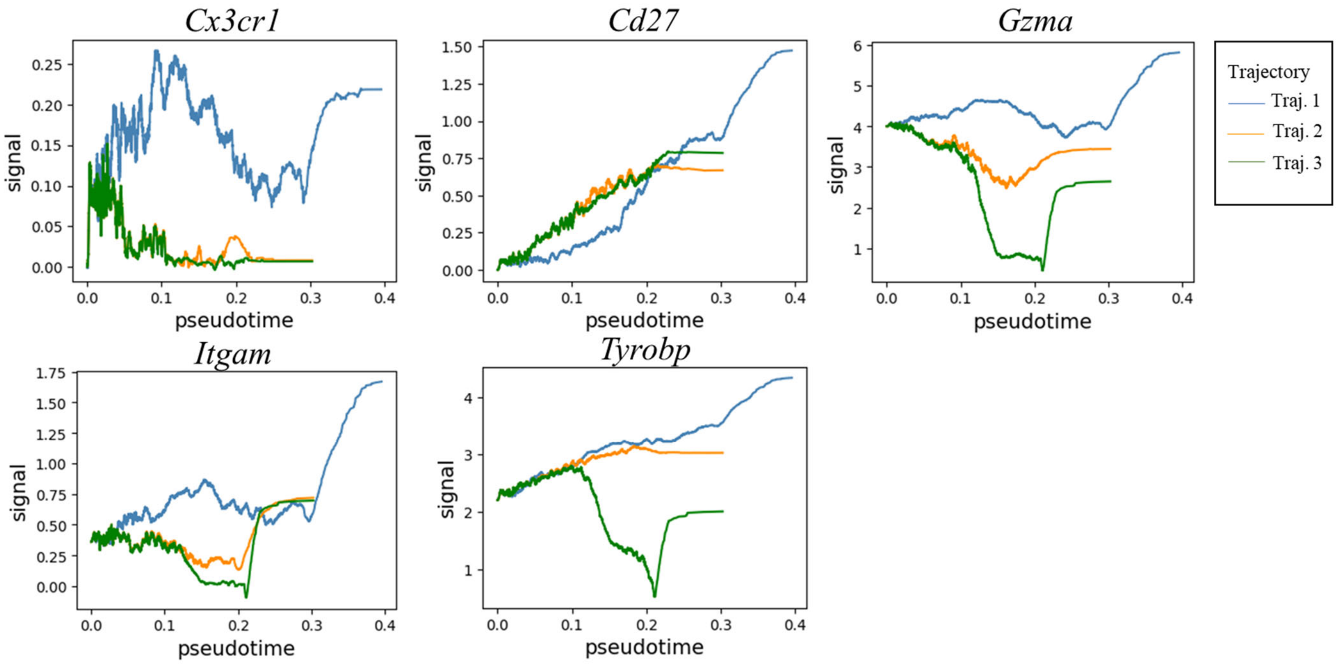


**Figure S 10** The signal curve of GO-BP map layer 2 GO terms from immune response cluster (GO-c1) across pseudotime. The blue curve corresponds to trajectory #1, the orange curve to trajectory #2, and the green curve to trajectory #3.

### Tables

**Table S 1** The -log(PMF) cutoff screening to maximize ARI, the product of the R-squared ($R^2_{degree}$.) of power-law distribution fit and the Aggregation Index (AI) of networks in GO categories. The red font indicates the best threshold of -log(PMF).

| **-log(PMF)** | **BP** | | | **MF** | | | **CC** | | |
|---|---|---|---|---|---|---|---|---|---|
| | **ARI** | $R^2_{degree}$ | **AI** | **ARI** | $R^2_{degree}$ | **AI** | **ARI** | $R^2_{degree}$ | **AI** |
| **0** | 0.838 | 0.84 | 1.00 | 0.710 | 0.71 | 1.00 | 0.774 | 0.77 | 1.00 |
| **1** | 0.844 | 0.84 | 1.00 | 0.746 | 0.75 | 1.00 | 0.794 | 0.79 | 1.00 |
| **2** | 0.847 | 0.85 | 1.00 | 0.805 | 0.80 | 1.00 | 0.836 | 0.84 | 1.00 |
| **3** | 0.863 | 0.86 | 1.00 | 0.841 | 0.84 | 1.00 | 0.853 | 0.85 | 1.00 |
| **4** | 0.871 | 0.87 | 1.00 | 0.857 | 0.86 | 1.00 | 0.853 | 0.85 | 1.00 |
| **5** | 0.875 | 0.88 | 1.00 | 0.843 | 0.85 | 1.00 | 0.867 | 0.87 | 0.99 |
| **6** | 0.883 | 0.88 | 1.00 | 0.836 | 0.85 | 0.99 | 0.878 | 0.89 | 0.99 |
| **7** | 0.891 | 0.89 | 1.00 | 0.823 | 0.85 | 0.97 | 0.852 | 0.88 | 0.97 |
| **8** | 0.887 | 0.89 | 0.99 | 0.808 | 0.86 | 0.94 | 0.853 | 0.90 | 0.95 |
| **9** | 0.884 | 0.90 | 0.98 | 0.782 | 0.86 | 0.91 | 0.814 | 0.89 | 0.92 |
| **10** | 0.860 | 0.89 | 0.96 | 0.750 | 0.87 | 0.86 | 0.796 | 0.89 | 0.89 |

**Table S 2** The α cutoff screening to maximize the BRI, the product of the $R^2_{layer}$ and baseline match score for generating GO layers. The red font indicates the best α threshold.

| α | BP | | | MF | | | CC | | |
|---|---|---|---|---|---|---|---|---|---|
| | **BRI** | $R^2_{layer}$ | **Baseline match** | **BRI** | $R^2_{layer}$ | **Baseline match** | **BRI** | $R^2_{layer}$ | **Baseline match** |
| **0** | 0.428 | 0.94 | 0.46 | 0.422 | 0.85 | 0.50 | 0.435 | 0.90 | 0.48 |
| **0.1** | 0.480 | 0.98 | 0.49 | 0.516 | 0.94 | 0.55 | 0.513 | 0.97 | 0.53 |
| **0.2** | 0.521 | 0.99 | 0.53 | 0.510 | 0.93 | 0.55 | 0.519 | 0.98 | 0.53 |
| **0.3** | 0.521 | 0.99 | 0.53 | 0.576 | 0.95 | 0.61 | 0.568 | 0.98 | 0.58 |
| **0.4** | 0.559 | 0.99 | 0.56 | 0.576 | 0.95 | 0.61 | 0.568 | 0.98 | 0.58 |
| **0.5** | 0.645 | 0.99 | 0.65 | 0.696 | 0.94 | 0.74 | 0.622 | 0.98 | 0.63 |
| **0.6** | 0.744 | 0.99 | 0.75 | 0.761 | 0.93 | 0.82 | 0.731 | 0.96 | 0.76 |
| **0.7** | 0.912 | 0.98 | 0.93 | 0.880 | 0.88 | 1.00 | 0.970 | 0.97 | 1.00 |
| **0.8** | 0.686 | 0.98 | 0.70 | 0.457 | 0.92 | 0.50 | 0.674 | 0.97 | 0.70 |
| **0.9** | 0.188 | 0.97 | 0.19 | 0.058 | 0.86 | 0.07 | 0.200 | 0.94 | 0.21 |

**Table S 3** The comparison of degree of inequality (DI) across the GO stratification approaches, GO depth, GO level and GO layer.

| | GO depth | GO level | GO layer |
|---|---|---|---|
| **BP** | -0.06 | -0.13 | 0.39 |
| **MF** | 0.12 | 0.08 | 0.57 |
| **CC** | -0.15 | -0.18 | 0.46 |

**Table S 4** Silhouette coefficients, Davies-Bouldin index, and Calinski-Harabasz index for selecting the optimal clustering algorithm and the optimal cluster number denoted by k.

| Method | category | cluster | k-means | k-medoids | hierarchy | spectrum |
|---|---|---|---|---|---|---|
| **Silhouette coefficient** | BP | 6 | 0.443 | 0.443 | 0.389 | 0.413 |
| | MF | 6 | 0.477 | 0.475 | 0.467 | 0.447 |
| | CC | 2 | 0.515 | 0.516 | 0.494 | 0.509 |
| **Davies-Bouldin index** | BP | 6 | 0.733 | 0.759 | 0.787 | 0.801 |
| | MF | 2 | 1.017 | 1.018 | 1.076 | 0.533 |
| | CC | 7 | 0.745 | 0.747 | 0.839 | 0.709 |
| **Calinski-Harabasz index** | BP | 8 | 1.8E+04 | 1.8E+04 | 1.6E+04 | 1.2E+04 |
| | MF | 6 | 7.1E+03 | 7.0E+03 | 6.8E+03 | 5.2E+03 |
| | CC | 7 | 3.1E+03 | 3.1E+03 | 2.6E+03 | 2.7E+03 |

**Table S 5** The GO-BP layers 2 and 3 GO terms from the immune response functional cluster enriched by the trajectory 1's DEGs in mNK cells.

| GO cluster | Layer | GO ID | GO name | Transition | Ovlp. | Genes | SSI | log(FC) |
|---|---|---|---|---|---|---|---|---|
| 1 | 2 | GO:0021626 | central nervous system maturation | 2_3 | 1/2 | CX3CR1 | 2.9 | 1.3 |
| | | | | 9_7 | 1/2 | CX3CR1 | -2.5 | -1.1 |
| | | GO:0150089 | multiple spine synapse organization | 2_3 | 1/1 | CX3CR1 | 3 | 1.3 |
| | | | | 9_7 | 1/1 | CX3CR1 | -2.7 | -1.1 |
| | | GO:0150090 | multiple spine synapse organization, single dendrite | 2_3 | 1/1 | CX3CR1 | 3 | 1.3 |
| | | | | 9_7 | 1/1 | CX3CR1 | -2.7 | -1.1 |
| | | GO:1904150 | negative regulation of microglial cell mediated cytotoxicity | 2_3 | 1/1 | CX3CR1 | 3 | 1.3 |
| | | | | 9_7 | 1/1 | CX3CR1 | -2.7 | -1.1 |
| | | GO:0090717 | adaptive immune memory response involving T cells and B cells | 3_9 | 1/3 | CD27 | 4.3 | 2 |
| | | | | 9_7 | 1/3 | CD27 | 4.3 | 2 |
| | 3 | GO:0033634 | positive regulation of cell-cell adhesion mediated by integrin | 3_9 | 1/6 | CD3E | 6.3 | 3.2 |
| | | | | 9_7 | 1/6 | CD3E | 6.1 | 3.1 |
| | | GO:0035705 | T-helper 17 cell chemotaxis | 3_9 | 1/1 | CCR2 | 4.7 | 1.9 |
| | | | | 9_7 | 1/1 | CCR2 | 4.1 | 1.6 |
| | | GO:0035782 | mature natural killer cell chemotaxis | 9_7 | 1/1 | XCL1 | 5.5 | 2.1 |
| | | GO:0043310 | negative regulation of eosinophil degranulation | 3_9 | 1/1 | CCR2 | 4.7 | 1.9 |
| | | | | 9_7 | 1/1 | CCR2 | 4.1 | 1.6 |
| | | GO:0071661 | regulation of granzyme B production | 9_7 | 1/2 | XCL1 | 5 | 2.1 |
| | | GO:0071663 | positive regulation of granzyme B production | 9_7 | 1/2 | XCL1 | 5 | 2.1 |
| | | GO:0090265 | positive regulation of immune complex clearance by monocytes and macrophages | 3_9 | 1/2 | CCR2 | 4.3 | 1.9 |
| | | | | 9_7 | 1/2 | CCR2 | 3.7 | 1.6 |
| | | GO:0090713 | immunological memory process | 3_9 | 1/6 | CD27 | 3.9 | 2 |
| | | | | 9_7 | 1/6 | CD27 | 3.9 | 2 |
| | | GO:0090716 | adaptive immune memory response | 3_9 | 1/4 | CD27 | 4.2 | 2 |
| | | | | 9_7 | 1/4 | CD27 | 4.1 | 2 |
| | | GO:0140507 | granzyme-mediated programmed cell death signaling pathway | 2_3 | 1/11 | GZMB | 2.1 | 1.1 |
| | | GO:2000409 | positive regulation of T cell extravasation | 3_9 | 1/4 | CCR2 | 3.9 | 1.9 |
| | | | | 9_7 | 1/4 | CCR2 | 3.3 | 1.6 |
| | | GO:2000458 | regulation of astrocyte chemotaxis | 3_9 | 1/2 | CCR2 | 4.3 | 1.9 |
| | | | | 9_7 | 1/2 | CCR2 | 3.7 | 1.6 |
| | | GO:2000464 | positive regulation of astrocyte chemotaxis | 3_9 | 1/1 | CCR2 | 4.7 | 1.9 |
| | | | | 9_7 | 1/1 | CCR2 | 4.1 | 1.6 |
| | | GO:2000511 | regulation of granzyme A production | 9_7 | 1/1 | XCL1 | 5.5 | 2.1 |
| | | GO:2000513 | positive regulation of granzyme A production | 9_7 | 1/1 | XCL1 | 5.5 | 2.1 |
| | | GO:2000517 | regulation of T-helper 1 cell activation | 9_7 | 1/1 | XCL1 | 5.5 | 2.1 |
| | | GO:2000518 | negative regulation of T-helper 1 cell activation | 9_7 | 1/1 | XCL1 | 5.5 | 2.1 |
| | | GO:2000537 | regulation of B cell chemotaxis | 9_7 | 1/2 | XCL1 | 5 | 2.1 |
| | | GO:2000538 | positive regulation of B cell chemotaxis | 9_7 | 1/2 | XCL1 | 5 | 2.1 |
| | | GO:2000557 | regulation of immunoglobulin production in mucosal tissue | 9_7 | 1/2 | XCL1 | 5 | 2.1 |
| | | GO:2000558 | positive regulation of immunoglobulin production in mucosal tissue | 9_7 | 1/2 | XCL1 | 5 | 2.1 |

**Table S 6** The GO-BP layers 2 and 3 GO terms from the immune response functional cluster enriched by the trajectory 2's DEGs in mNK cells.

| GO Cluster | Layer | GO ID | GO name | Transition | Ovlp. | Genes | Score | LFC |
|---|---|---|---|---|---|---|---|---|
| 1 | 2 | GO:0021626 | central nervous system maturation | 1_0 | 1/2 | CX3CR1 | -4.8 | -2.1 |
| | | GO:0090717 | adaptive immune memory response involving T cells and B cells | 1_0 | 1/3 | CD27 | 4.2 | 2 |
| | | GO:0141201 | pyroptotic cell death | 0_4 | 1/5 | GZMA | -2.1 | -1.1 |
| | | GO:0150089 | multiple spine synapse organization | 1_0 | 1/1 | CX3CR1 | -5.2 | -2.1 |
| | | GO:0150090 | multiple spine synapse organization, single dendrite | 1_0 | 1/1 | CX3CR1 | -5.2 | -2.1 |
| | | GO:1902483 | cytotoxic T cell pyroptotic cell death | 0_4 | 1/2 | GZMA | -2.3 | -1.1 |
| | | GO:1904150 | negative regulation of microglial cell mediated cytotoxicity | 1_0 | 1/1 | CX3CR1 | -5.2 | -2.1 |
| | | GO:1904643 | response to curcumin | 0_4 | 1/1 | ITGAM | -2.2 | -1 |
| | 3 | GO:0035705 | T-helper 17 cell chemotaxis | 1_0 | 1/1 | CCR2 | 4.9 | 2 |
| | | | | 2_1 | 1/1 | CCR2 | 2.6 | 1.3 |
| | | GO:0035782 | mature natural killer cell chemotaxis | 0_4 | 1/1 | XCL1 | 3.1 | 1.4 |
| | | | | 1_0 | 1/1 | XCL1 | 4.7 | 1.9 |
| | | GO:0043310 | negative regulation of eosinophil degranulation | 1_0 | 1/1 | CCR2 | 4.9 | 2 |
| | | | | 2_1 | 1/1 | CCR2 | 2.6 | 1.3 |
| | | GO:0045914 | negative regulation of catecholamine metabolic process | 0_4 | 1/4 | ITGAM | -1.9 | -1 |
| | | GO:0045963 | negative regulation of dopamine metabolic process | 0_4 | 1/4 | ITGAM | -1.9 | -1 |
| | | GO:0071661 | regulation of granzyme B production | 0_4 | 1/2 | XCL1 | 2.9 | 1.4 |
| | | | | 1_0 | 1/2 | XCL1 | 4.3 | 1.9 |
| | | GO:0071663 | positive regulation of granzyme B production | 0_4 | 1/2 | XCL1 | 2.9 | 1.4 |
| | | | | 1_0 | 1/2 | XCL1 | 4.3 | 1.9 |
| | | GO:0090265 | positive regulation of immune complex clearance by monocytes and macrophages | 1_0 | 1/2 | CCR2 | 4.5 | 2 |
| | | | | 2_1 | 1/2 | CCR2 | 2.6 | 1.3 |
| | | GO:0090713 | immunological memory process | 1_0 | 1/6 | CD27 | 3.8 | 2 |
| | | GO:0090716 | adaptive immune memory response | 1_0 | 1/4 | CD27 | 4 | 2 |
| | | GO:0140507 | granzyme-mediated programmed cell death signaling pathway | 0_4 | 1/11 | GZMA | -1.9 | -1.1 |
| | | | | 1_0 | 1/11 | GZMB | -1.9 | -1.1 |
| | | GO:0150064 | vertebrate eye-specific patterning | 0_4 | 1/3 | ITGAM | -2 | -1 |
| | | GO:2000409 | positive regulation of T cell extravasation | 1_0 | 1/4 | CCR2 | 4.1 | 2 |
| | | | | 2_1 | 1/4 | CCR2 | 2.6 | 1.3 |
| | | GO:2000458 | regulation of astrocyte chemotaxis | 1_0 | 1/2 | CCR2 | 4.5 | 2 |
| | | | | 2_1 | 1/2 | CCR2 | 2.6 | 1.3 |
| | | GO:2000464 | positive regulation of astrocyte chemotaxis | 1_0 | 1/1 | CCR2 | 4.9 | 2 |
| | | | | 2_1 | 1/1 | CCR2 | 2.6 | 1.3 |
| | | GO:2000511 | regulation of granzyme A production | 0_4 | 1/1 | XCL1 | 3.1 | 1.4 |
| | | | | 1_0 | 1/1 | XCL1 | 4.7 | 1.9 |
| | | GO:2000513 | positive regulation of granzyme A production | 0_4 | 1/1 | XCL1 | 3.1 | 1.4 |
| | | | | 1_0 | 1/1 | XCL1 | 4.7 | 1.9 |
| | | GO:2000517 | regulation of T-helper 1 cell activation | 0_4 | 1/1 | XCL1 | 3.1 | 1.4 |
| | | | | 1_0 | 1/1 | XCL1 | 4.7 | 1.9 |
| | | GO:2000518 | negative regulation of T-helper 1 cell activation | 0_4 | 1/1 | XCL1 | 3.1 | 1.4 |
| | | | | 1_0 | 1/1 | XCL1 | 4.7 | 1.9 |
| | | GO:2000537 | regulation of B cell chemotaxis | 0_4 | 1/2 | XCL1 | 2.9 | 1.4 |
| | | | | 1_0 | 1/2 | XCL1 | 4.3 | 1.9 |
| | | GO:2000538 | positive regulation of B cell chemotaxis | 0_4 | 1/2 | XCL1 | 2.9 | 1.4 |

| | | | | | | | | |
|---|---|---|---|---|---|---|---|---|
| | | | | 1_0 | 1/2 | XCL1 | 4.3 | 1.9 |
| | | GO:2000557 | regulation of immunoglobulin production in mucosal tissue | 0_4 | 1/2 | XCL1 | 2.9 | 1.4 |
| | | | | 1_0 | 1/2 | XCL1 | 4.3 | 1.9 |
| | | GO:2000558 | positive regulation of immunoglobulin production in mucosal tissue | 0_4 | 1/2 | XCL1 | 2.9 | 1.4 |
| | | | | 1_0 | 1/2 | XCL1 | 4.3 | 1.9 |

**Table S 7** The GO-BP layers 2 and 3 GO terms from the immune response functional cluster enriched by the trajectory 3's DEGs in mNK cells.

| GO Cluster | Layer | GO ID | GO name | Path | Ovlp. | Genes | Score | LFC |
|---|---|---|---|---|---|---|---|---|
| 1 | 2 | GO:0002222 | stimulatory killer cell immunoglobulin-like receptor signaling pathway | 0_12 | 1/2 | TYROBP | -2.4 | -1 |
| | | | | 12_8 | 1/2 | TYROBP | -4.3 | -2 |
| | | GO:0021626 | central nervous system maturation | 1_0 | 1/2 | CX3CR1 | -4.8 | -2 |
| | | GO:0090717 | adaptive immune memory response involving T cells and B cells | 1_0 | 1/3 | CD27 | 4.2 | 2 |
| | | GO:0141201 | pyroptotic cell death | 0_12 | 1/5 | GZMA | -8.8 | -5 |
| | | GO:0150089 | multiple spine synapse organization | 1_0 | 1/1 | CX3CR1 | -5.2 | -2 |
| | | GO:0150090 | multiple spine synapse organization, single dendrite | 1_0 | 1/1 | CX3CR1 | -5.2 | -2 |
| | | GO:1902483 | cytotoxic T cell pyroptotic cell death | 0_12 | 1/2 | GZMA | -9.7 | -5 |
| | | GO:1904150 | negative regulation of microglial cell mediated cytotoxicity | 1_0 | 1/1 | CX3CR1 | -5.2 | -2 |
| | | GO:1904643 | response to curcumin | 0_12 | 1/1 | ITGAM | -8.6 | -4 |
| | 3 | GO:0002418 | immune response to tumor cell | 0_12 | 1/7 | PRF1 | -4.3 | -2 |
| | | GO:0033634 | positive regulation of cell-cell adhesion mediated by integrin | 12_8 | 1/6 | CD3E | 15.3 | 7.1 |
| | | GO:0035705 | T-helper 17 cell chemotaxis | 12_8 | 1/1 | CCR2 | -5.7 | -2 |
| | | | | 1_0 | 1/1 | CCR2 | 4.9 | 2 |
| | | | | 2_1 | 1/1 | CCR2 | 2.6 | 1.3 |
| | | GO:0035782 | mature natural killer cell chemotaxis | 1_0 | 1/1 | XCL1 | 4.7 | 1.9 |
| | | | | 0_12 | 1/1 | XCL1 | 5 | 2.2 |
| | | | | 12_8 | 1/1 | XCL1 | -4.5 | -2 |
| | | GO:0042590 | antigen processing and presentation of exogenous peptide antigen via MHC class I | 12_8 | 1/9 | FCER1G | -3.4 | -2 |
| | | GO:0043310 | negative regulation of eosinophil degranulation | 12_8 | 1/1 | CCR2 | -5.7 | -2 |
| | | | | 1_0 | 1/1 | CCR2 | 4.9 | 2 |
| | | | | 2_1 | 1/1 | CCR2 | 2.6 | 1.3 |
| | | GO:0045914 | negative regulation of catecholamine metabolic process | 0_12 | 1/4 | ITGAM | -7.3 | -4 |
| | | GO:0045963 | negative regulation of dopamine metabolic process | 0_12 | 1/4 | ITGAM | -7.3 | -4 |
| | | GO:0051712 | positive regulation of killing of cells of another organism | 0_12 | 1/4 | PRF1 | -4.6 | -2 |
| | | GO:0071661 | regulation of granzyme B production | 1_0 | 1/2 | XCL1 | 4.3 | 1.9 |
| | | | | 0_12 | 1/2 | XCL1 | 4.6 | 2.2 |
| | | | | 12_8 | 1/2 | XCL1 | -4.1 | -2 |
| | | GO:0071663 | positive regulation of granzyme B production | 0_12 | 1/2 | XCL1 | 4.6 | 2.2 |
| | | | | 12_8 | 1/2 | XCL1 | -4.1 | -2 |
| | | | | 1_0 | 1/2 | XCL1 | 4.3 | 1.9 |
| | | GO:0090265 | positive regulation of immune complex clearance by monocytes and macrophages | 12_8 | 1/2 | CCR2 | -5.2 | -2 |
| | | | | 1_0 | 1/2 | CCR2 | 4.5 | 2 |
| | | | | 2_1 | 1/2 | CCR2 | 2.6 | 1.3 |
| | | GO:0090713 | immunological memory process | 1_0 | 1/6 | CD27 | 3.8 | 2 |
| | | GO:0090716 | adaptive immune memory response | 1_0 | 1/4 | CD27 | 4 | 2 |
| | | GO:0140507 | granzyme-mediated programmed cell death signaling pathway | 0_12 | 3/11 | PRF1;GZMA;GZMB | -17 | -3 |
| | | | | 1_0 | 1/11 | GZMB | -1.9 | -1 |
| | | GO:0150064 | vertebrate eye-specific patterning | 0_12 | 1/3 | ITGAM | -7.6 | -4 |
| | | GO:1902685 | positive regulation of receptor localization to synapse | 0_12 | 1/4 | TYROBP | -2.2 | -1 |
| | | | | 12_8 | 1/4 | TYROBP | -3.9 | -2 |

| | | | | | | | | |
|---|---|---|---|---|---|---|---|---|
| | | GO:2000409 | positive regulation of T cell extravasation | 12_8 | 1/4 | CCR2 | -4.7 | -2 |
| | | | | 1_0 | 1/4 | CCR2 | 4.1 | 2 |
| | | | | 2_1 | 1/4 | CCR2 | 2.6 | 1.3 |
| | | GO:2000458 | regulation of astrocyte chemotaxis | 12_8 | 1/2 | CCR2 | -5.2 | -2 |
| | | | | 1_0 | 1/2 | CCR2 | 4.5 | 2 |
| | | | | 2_1 | 1/2 | CCR2 | 2.6 | 1.3 |
| | | GO:2000464 | positive regulation of astrocyte chemotaxis | 12_8 | 1/1 | CCR2 | -5.7 | -2 |
| | | | | 1_0 | 1/1 | CCR2 | 4.9 | 2 |
| | | | | 2_1 | 1/1 | CCR2 | 2.6 | 1.3 |
| | | GO:2000511 | regulation of granzyme A production | 0_12 | 1/1 | XCL1 | 5 | 2.2 |
| | | | | 12_8 | 1/1 | XCL1 | -4.5 | -2 |
| | | | | 1_0 | 1/1 | XCL1 | 4.7 | 1.9 |
| | | GO:2000513 | positive regulation of granzyme A production | 0_12 | 1/1 | XCL1 | 5 | 2.2 |
| | | | | 12_8 | 1/1 | XCL1 | -4.5 | -2 |
| | | | | 1_0 | 1/1 | XCL1 | 4.7 | 1.9 |
| | | GO:2000517 | regulation of T-helper 1 cell activation | 0_12 | 1/1 | XCL1 | 5 | 2.2 |
| | | | | 12_8 | 1/1 | XCL1 | -4.5 | -2 |
| | | | | 1_0 | 1/1 | XCL1 | 4.7 | 1.9 |
| | | GO:2000518 | negative regulation of T-helper 1 cell activation | 0_12 | 1/1 | XCL1 | 5 | 2.2 |
| | | | | 12_8 | 1/1 | XCL1 | -4.5 | -2 |
| | | | | 1_0 | 1/1 | XCL1 | 4.7 | 1.9 |
| | | GO:2000537 | regulation of B cell chemotaxis | 0_12 | 1/2 | XCL1 | 4.6 | 2.2 |
| | | | | 12_8 | 1/2 | XCL1 | -4.1 | -2 |
| | | | | 1_0 | 1/2 | XCL1 | 4.3 | 1.9 |
| | | GO:2000538 | positive regulation of B cell chemotaxis | 0_12 | 1/2 | XCL1 | 4.6 | 2.2 |
| | | | | 12_8 | 1/2 | XCL1 | -4.1 | -2 |
| | | | | 1_0 | 1/2 | XCL1 | 4.3 | 1.9 |
| | | GO:2000557 | regulation of immunoglobulin production in mucosal tissue | 0_12 | 1/2 | XCL1 | 4.6 | 2.2 |
| | | | | 12_8 | 1/2 | XCL1 | -4.1 | -2 |
| | | | | 1_0 | 1/2 | XCL1 | 4.3 | 1.9 |
| | | GO:2000558 | positive regulation of immunoglobulin production in mucosal tissue | 0_12 | 1/2 | XCL1 | 4.6 | 2.2 |
| | | | | 12_8 | 1/2 | XCL1 | -4.1 | -2 |
| | | | | 1_0 | 1/2 | XCL1 | 4.3 | 1.9 |
| | | GO:2001206 | positive regulation of osteoclast development | 0_12 | 1/4 | TYROBP | -2.2 | -1 |
| | | | | 12_8 | 1/4 | TYROBP | -3.9 | -2 |
